# Pig manure treatment strategies for mitigating the spread of antibiotic resistance

**DOI:** 10.1101/2022.05.17.492273

**Authors:** Magdalena Zalewska, Aleksandra Błażejewska, Agnieszka Czapko, Magdalena Popowska

## Abstract

One of the most important public health challenges facing the world today is that posed by antibiotic resistance. Many pathogenic antibiotic-resistant bacteria and their antibiotic resistance genes, usually located on mobile genetic elements, are frequently present in the faeces of farm animals. To prevent the possibility of antimicrobial resistance transfer to the environment, these faeces should undergo treatment before being used as natural fertilizer. The two strategies for processing pig manure proposed in this study, *viz.* storage (most commonly used for livestock manure today) and composting, are cheap and do not require special tools or technologies. The present study examines the changes in the physicochemical properties of treated manure, in the microbiome, through metagenomic sequencing, and in the resistome, using the SmartChip Real-time PCR system compared to raw manure. This is the first such comprehensive analysis performed on the same batch of manure. Our results suggest that while none of the processes completely eliminates the environmental risk, composting results in a faster and more pronounced reduction of mobile genetic elements harbouring antibiotic resistance genes, including those responsible for multi-drug resistance. The physicochemical parameters of the treated manure are comparable after both processes; however, composting resulted in significantly higher organic matter. Overall, it appears that the composting process can be an efficient strategy for mitigating the spread of antibiotic resistance in the environment and reducing the risk of its transfer to agricultural crops and hence, the food chain. It also provides the organic matter necessary for humus formation, and increases the sorption properties of the soil and the micro and macro elements necessary for plant growth, which in turn translates into increased soil productivity.

**Highlights:**

- The changes in microbial population composition correlate with changes in specific antibiotic-resistance genes and mobile genetic elements in the studied populations.
- Positive correlations have been demonstrated between microbial phyla and genes coding the multi-drug resistance mechanism
- Co-occurrence networks showed positive correlations between antibiotic-resistance genes and mobile genetic elements
- The composting strategy was most efficient at reducing microbial loads, antibiotic resistance genes and mobile genetic elements.
- Composted manure can be part of a natural, safe soil fertilization strategy.

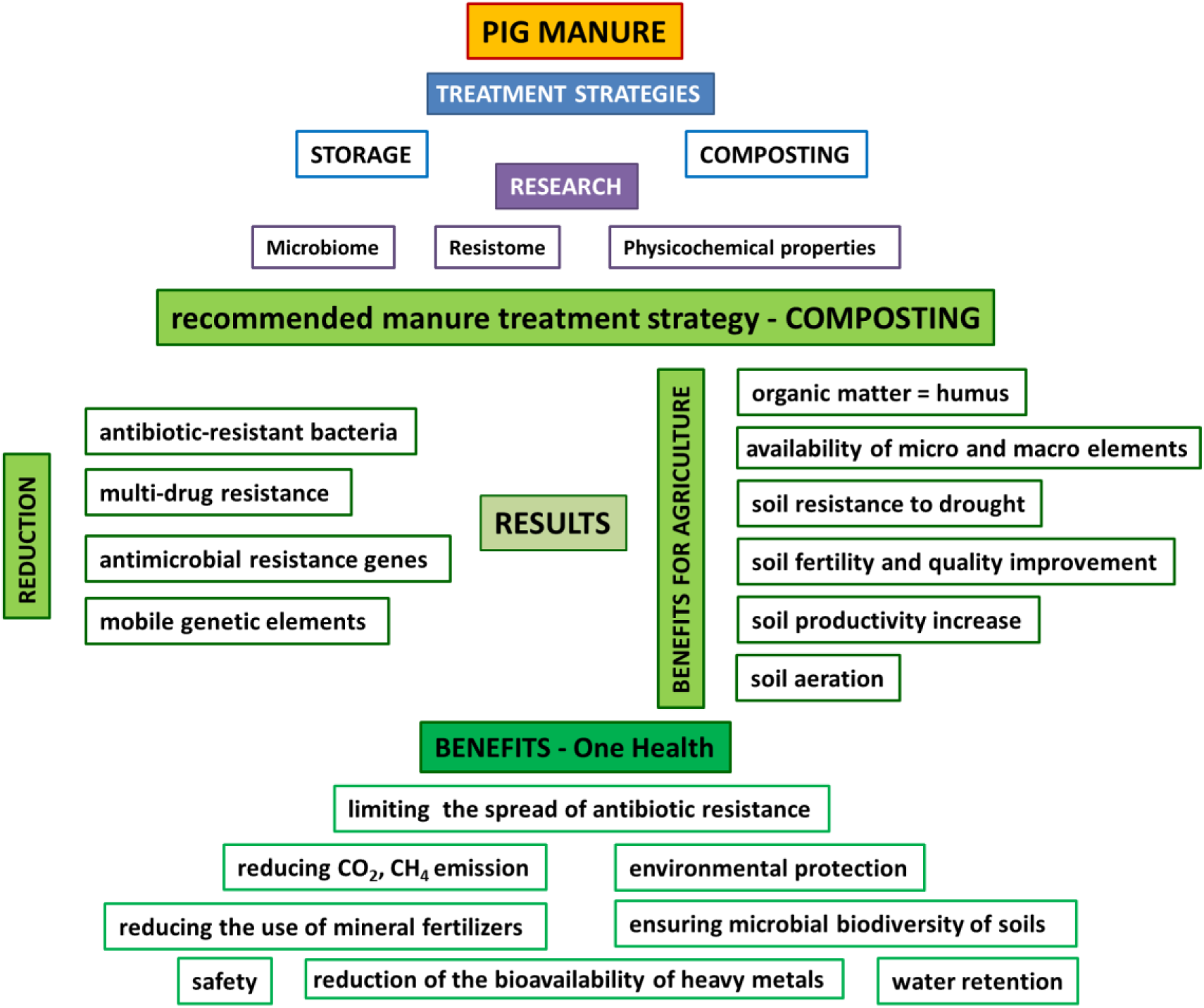

## 1. Introduction

The growing demand for meat has led to an increase in pig farming worldwide. Annual global pig production has increased in the last 50 years, reaching approximately 120 million tons in 2018. Total swine meat consumption is estimated to rise by 13% in 2030 and 22% in 2050 compared with 2020 (Ritchie and Roser, 2017). The main pig meat producers are China (45% of global production), followed by the United States, Germany, Spain and Vietnam (FAO, 2020), with these five countries accounting for nearly 65% of world pig meat production. In the European Union, approximately 30% of pig farming is concentrated in regions ranging from Denmark through north-western Germany and the Netherlands to northern Belgium, with Cataluña and Murcia in Spain, Lombardy in Italy, and Bretagne in France also being important regions (Makara and Kowalski, 2018). While in the EU, antibiotics are only applied to treat bacterial infections, in some other countries, they are also commonly applied to promote growth and increase production efficiency; in these countries, approximately 66,667 tons of antibiotics are applied to livestock worldwide each year (Menz et al., 2019). However, due to the link reported between antibiotic use in livestock and the emergence of antibiotic resistance in pathogenic bacteria, the use of antibiotics as growth promoters is being limited and has been prohibited in the EU since 2006 (Dibner and Richards, 2005); it has also been restricted in the United States since 2017 (Centner, 2016) and in China since 2020 (Hu and Cowling, 2020). Nevertheless, China remains the largest producer and consumer of antibiotics, with 52% being used in animal production (Wang et al., 2021).

The choice of antibiotic compound depends on the animal production sector. Swine typically receive more antibiotics than cattle because of their dense breeding environment and higher exposure to bacterial diseases. They are most commonly treated with tetracyclines (predominantly oxytetracycline and doxycycline), macrolides (predominantly tylosin), and sulfonamides (predominantly sulfadiazine and sulfamethoxazole) combined with trimethoprim (Berendsen et al., 2018). While the swine gastrointestinal microbiota harbours a diverse population of bacteria known to support the health of the host, they may also be a source of drug resistance genes. Intensive livestock farms are characterized by a combination of high bacterial load and high antimicrobial selection pressure: conditions known to promote the occurrence of antimicrobial-resistant bacteria (ARB) and antibiotic resistance genes (ARGs) (Luiken et al., 2020). In addition, disturbances in gut microbiota may enhance ARG transfer or enhance the abundance of ARB shed from the animal.

Post-weaning, piglets are highly susceptible to many diseases, and antibiotics are commonly added to food or water as prophylactic treatments. As a result of such treatment, the swine gastrointestinal microbiota undergoes shifts of varying lengths, with the resulting changes in taxa depending on the antibiotic and intestinal segment. It has also been demonstrated that oral administration alters the porcine intestinal microbiome more than injections (Ricker et al., 2020).

Typically, more than half the administered dose of antibiotics is excreted unchanged in the urine and faeces. Only a small number of antibiotics are partially metabolized by the host animal, yielding microbiologically active or inactive metabolites. For example, in the liver, enrofloxacin is partially (<25%) metabolized to ciprofloxacin, while sulfonamides are metabolized to a small extent to the less active N4-acetylosulfonamides; in both cases, the products are microbiologically active (Anderson et al., 2012).

As a single pig may produce up to 6.4 kg of wet manure per day (Girotto and Cossu, 2017), large animal breeding facilities produce vast amounts of animal faeces, resulting in the production of 1.7 billion tons of faeces annually worldwide (Y. Zhang et al., 2021). In pig farms in Germany, Spain, the UK and the Netherlands alone, annual manure production amounts to over 120 million tons (Makara and Kowalski, 2018). The easiest and cheapest way to dispose of such waste is land application; however, livestock manure has been identified as a reservoir of antibiotics, ARGs, and potentially pathogenic ARB, posing a considerable threat to animals and human health. Indeed, pig manure is regarded as a key source of the spread of antibiotic resistance to the environment (Lu and Lu, 2019; Zalewska et al., 2021). Certainly, the direct application of livestock manure originating from farm animals treated with antibiotics is known to expose agricultural areas to high levels of antibiotics. Depending on the physio-chemical properties of the antibiotic and soil composition, the compounds may remain in the manure-amended soil, or may be transferred to ground and surface waters by leaching or runoff. In addition, the rate of degradation varies depending on the antibiotic type, and in many cases, this can be very low for up to 30 days (Zalewska et al., 2021). When in soil, the compounds may be taken up by crops, where they accumulate and then enter the food chain. In addition, the constant presence of a low concentration of antibiotics and their active residues in the environment favours the selection of resistant bacteria (Zalewska et al., 2021).

ARGs, are compounds whose presence in the environment can be more harmful than the antibiotic itself (Larrañaga et al., 2018). A recent study found the most prevalent classes of ARGs in French and Danish swine to be tetracycline-, beta-lactam-, macrolide-, streptogramin-, and bacitracin-resistance genes; in contrast, the ARG profile in Chinese pigs was found to be dominated by tetracycline-, aminoglycoside-, and beta-lactam resistance genes, which was consistent with the antibiotic use profile (Xiao et al., 2016; Zhu et al., 2013). The abundance of ARGs is, in general, higher in swine than in cattle. A study of the antimicrobial susceptibility of *E. coli* isolated from swine faeces found resistance rates of 66.7% for tetracyclines, 66.7% for aminoglycosides, 66.7% for sulfonamides, and 33.3% for quinolones. In addition, the isolates showed resistance against almost all antibiotic types used for swine (Lim et al., 2020).

The spread of ARGs in the environment is a cyclical process based on vertical gene transfer (VGT), through the succession of host bacteria, and horizontal gene transfer (HGT), through the transmission of ARGs via mobile genetic elements (MGEs) (Jia et al., 2017; Wellington et al., 2013). HGT can accelerate the spread of ARGs in the environment (Piotrowska and Popowska, 2014). Interestingly, the presence of any heavy metals in the environment promotes co-selection and acts as a synergistic selection factor for ARGs (Zhu et al., 2013).

The most commonly-applied manure management strategy is storage, where the faeces are simply set aside in a designated secure location for a prolonged time. This requires sufficient storage capacity, as the minimal storage periods before spreading varies from 4 to 7.5 months, depending on the animal type, the length of time at pasture, and the geographical location (Loyon, 2018). Alternatively, manure can also be composted: an environmentally and economically-beneficial practice for processing solid organic waste, allowing for efficient volume reduction, pathogen removal, and antibiotic content reduction (Tasho and Cho, 2016). The most important factors include temperature, moisture, carbon to nitrogen ratio, aeration rate, pH level, and the physical structure of the raw material (Li et al., 2008). Composting is a biological process in which organic waste is transformed into a homogenous and plant-available material. It comprises a complex set of metabolic processes performed by different microorganisms using nitrogen and carbon to produce their biomass while generating heat (Azim et al., 2018). The process takes place under aerobic conditions, with adequate moisture and temperature, and is characterized by a long exothermic phase, during which the temperature of the composted substance increases to 65-78 °C. The final product of the process is stable, valuable, sanitary, safe natural fertilizer, and offers better quality content than stored manure (Boniecki et al., 2013). Furthermore, soil fertilized with natural fertilizers has a higher organic matter and total nitrogen content than soil fertilized chemically, and demonstrates greater productivity (Muscolo et al., 2018).

Despite this, the treatments used for manure before land spreading varies regionally, and, despite the clear advantages of composting, it is most commonly subjected to direct application; for example, in the Netherlands, untreated pig manure is widely applied on fields (85%) (Berendsen et al., 2018), while in France, manure management is mainly based on storage in buildings and manure pits, and direct spreading; manure treatment is not widely reported (Loyon, 2018). Generally, it is estimated that, in Europe, less than 10% of the total manure produced is treated by a single technology. Solid manure treatment strategies such as composting, drying, and combustion are applied to 0.8% of livestock manure production in the European Union. They are mostly used in Spain, where 3% of the manure production in the country is processed with these techniques (Herrero et al., 2015).

In Polish agriculture, the basic method of manure management is its use in direct fertilization of fields. The Institute of Plant Cultivation, Fertilization and Soil Science, estimates the total annual production of solid manure in Poland to be approximately 80 million tons, and approximately 21.5 million m^3^ as slurry. The use and storage of natural fertilizers (solid manure and slurry) are regulated by the Act on Fertilizers and Fertilization (10 July 2007) and Article 47 of the Water Law (18 July 2001), In addition, the maximum allowable doses of natural fertilizers is specified by the Nitrates Directive: the amount of nitrogen used in natural fertilizers cannot exceed 170 kg per year per hectare, and natural fertilizers can be used only from 1st March to 30th November (Herrero et al., 2015).

Currently, little data exists on the distribution of ARGs in pig manure, and its microbial community structure; in addition, no studies have compared the composting and storage of material obtained from the same source. The aim of the present study is to evaluate the role of pig manure as a crucial hot-spot for the spread of ARGs into the environment, even considering the strict control of antibiotic application for livestock in EU member countries. The findings highlight the need to introduce composting for treating manure before its land application, rather than its storage. It also determines the transmission potential of ARGs by identifying the correlations between microbial phyla and ARG groups, and between ARGs and MGEs observed during both storage and composting.

To determine how the spread of AMR can best be reduced before manure is applied to the fields, the study compares the effects of two commonly-used manure management strategies, storage and composting, at the laboratory scale. It is important to note that all samples used in the two methods were collected from the same animals from a single farm and at the same time point. The analysis consists mainly of high-throughput qPCR to determine the dynamics of ARG distribution, and sequencing protocols targeting V3-V4 variable regions of 16S rRNA to allow microbial community identification.

## 2. Materials and Method

### 2.1. Sample collection

The pig manure sample were collected from a Polish commercial finisher farm in Spring (April) 2019. Manure and slurry were separated on-farm. A solid fraction was collected in two blue plastic drums of approximately 20 L each (one for composting and the other for storage), and was taken to the laboratory before further processing and analysis. The farm owner agreed to manure sampling and shared the history of the use of antibiotics on the farm. The pig herd consists of 200 pigs (50 pigs per group pen), weight gain of 1100 g/day/pig. Diet used: 80% grain meal from own farm, 20% complementary feed: protein base, soybean meal or rapeseed meal + vitamins, micro- and macronutrients.

### 2.2. Pig manure management strategies and sampling

5kg of pig manure was mixed with Sitka spruce sawdust at the ratio of 4 (manure):1 (sawdust) (w/w) and placed in a bucket. Sawdust was obtained from the local hardware store. The organic matter, total nitrogen, phosphorus, calcium, magnesium, and heavy metal (Cr, Cu, Cd, Ni, Pb, Zn, Hg) content in raw manure and after storage (4M) and composting (10W) was determined.

#### 2.2.1. Composting

The container with the mixture was placed in the laboratory under the ventilated hood at ambient temperature. The mixture was turned (mixed well) each week for ten weeks. The temperature, moisture, and pH were measured in the compost mixture at about 15 cm (half the mixture’s height) each week. The moisture content was measured periodically and adjusted to 50-60% by adding sterile MilliQ water (to avoid potential contamination with ARGs or bacteria). The experiment was conducted in triplicate. Samples for the microbiological and molecular analysis were collected at the beginning, in the middle (fifth week), and at the end of the process (tenth week).

#### 2.2.2. Storage

The container with the mixture was then placed in the laboratory under the ventilated hood at ambient temperature; however, the sample was not mixed. The temperature, moisture, and pH were measured in the compost mixture at about 15 cm (half the mixture’s height) each week. The experiment was conducted in triplicate. Samples for the microbiological and molecular analysis were collected at the beginning of the process, in the middle (second month), and at the end (the fourth month). A four-month storage period was used, in accordance with EU legislation.

### 2.3. DNA extraction and metagenomic sequencing

Total DNA was isolated from 500 mg of the manure sample using FastDNA™ SPIN Kit for Feces (MP Biomedicals) following the manufacturer’s ’typical workflow’. The quality and quantity of extracted DNA were analysed with an Invitrogen Qubit 4.0 Fluorometer (dsDNA high-sensitivity assay kit) and Colibri spectrophotometer (Titertek Berthold). The DNA samples were isolated in triplicate and then pooled to obtain a single DNA sample for each time point.

The characteristics of the bacterial community structure were determined *via* 16S rRNA sequencing targeting the variable V3-V4 regions of bacterial 16S rRNA. Following the manufacturer’s recommendations, the libraries were prepared with Nextera® XT index Kit v2 Set A (Illumina). The prepared amplicons were then subjected to high-throughput sequencing using the MiSeq Illumina platform to generate 2x300 bp paired-end sequences at the DNA Sequencing and Oligonucleotide Synthesis Facility (Institute of Biochemistry and Biophysics Polish Academy of Sciences). Raw sequences were processed and analysed using Qiime2 software (Bolyen et al., 2019) with the DADA2 option for sequence quality control and the newest release of the SILVA ribosomal RNA sequence database for taxonomy assignment (Quast et al., 2013; Yilmaz et al., 2014). The obtained data was analysed and visualized using MicrobiomeAnalyst (Chong et al., 2020).

### 2.4. High-Throughput qPCR and Primers

The qPCR array was performed in the SmartChip Real-time PCR system by Resistomap. The qPCR cycling conditions and initial data processing were performed as described previously (Wang et al., 2014). The same 384-well template of primers was used for each DNA sample. Three technical replicates were run for each sample. The samples used for the analysis have similar quality: 260/280 ratio of 1.8-2.0 (+/- 0.1) and a concentration of 10 ng/μl. The concentration of the sample was measured using the Qubit 4.0 Fluorimeter (Thermo Fisher Scientific), and the quality using a Colibri spectrophotometer (Titertek Berthold). Only samples matching the described criteria were analysed. After amplification based on the qPCR SmartChip Real-Time PCR cycler (Takara), threshold cycle (CT) values were calculated using default parameters provided with the SmartChip analysis software; the genes and primer sequences are listed in Supplementary materials file 1. The gene names and groups listed and categorized in the table from Supplementary materials file 1 are used throughout the manuscript.

Melting curve and qPCR efficiency analysis were performed on all samples for each primer set. Amplicons with unspecific melting curves and multiple slope profile peaks were considered false positives and discarded from the analysis. Samples fitting the following criteria were taken for further analysis: (1) Ct values ≤ 27, (2) at least two replicates, (3) amplification efficiency in the range of 1.8 to 2.2. The relative copy number was calculated using equation 1 (Chen et al., 2016). The gene copy numbers were calculated by normalizing the relative copy numbers per 16S rRNA gene copy numbers.

Eq.1: Relative gene copy number = 10^(27-Ct)/(10/3)^

The relationships between ARGs and MGEs and between ARGs and microbial taxa were determined using Spearman’s correlation coefficient. The calculation of Spearman’s correlation coefficient and data visualization were performed with GraphPad Prism 9. The analysed relationships were used to construct networks using Cytoscape. The correlations were considered strong and significant when the absolute value of Spearman’s rank |r|>0.7 and p<0.05 (Zhu et al., 2017).

## 3. Results

### 3.1. Physio-chemical properties of pig manure

To ensure safe and sustainable management of pig manure, it is important to assess its physio-chemical properties. Therefore, the samples were studied for changes in heavy metal concentration during processing by measuring Chrome (Cr), Copper (Cu), Cadmium (Cd), Nickel (Ni), Lead (Pb), Zinc (Zn) and Mercury (Hg) at baseline and at the end of treatment. In addition, the initial and treated samples were also tested for pH, and microelement content, as well as the levels of the main crop nutrients (N, P), elements (Ca, Mg) and dry matter (DM) and organic matter content (OM) (Table S1).

The organic matter (OM) content was found to increase during both treatment strategies, but a greater increase was observed after composting. In addition, both sets of treated matter showed enrichment of N but a reduction in concentration of P, Mg and Ca compared with baseline. The manure pH was found to rise slightly from 7.0 to 7.4 during composting; however, a similar rise was observed during storage. While the concentrations of heavy metals also slightly increased after both processes, the concentration was remained safely below the applicable standards in Poland. The compost was found to be characterised by greater heavy metal contents than the stored manure, with Zn and Cu predominating.

### 3.1. Microbial composition of samples

The microbial composition was determined for the untreated pig manure samples, i.e. at baseline, and those subjected to composting and storage. In each case, the mean results from three replicates are presented. The microbial profiles of the bacterial community were obtained through 16S rRNA amplicon sequence analysis. The composition of bacterial communities at the phylum level is presented using the relative abundance graph type (Fig. 1).

**Figure 1:**
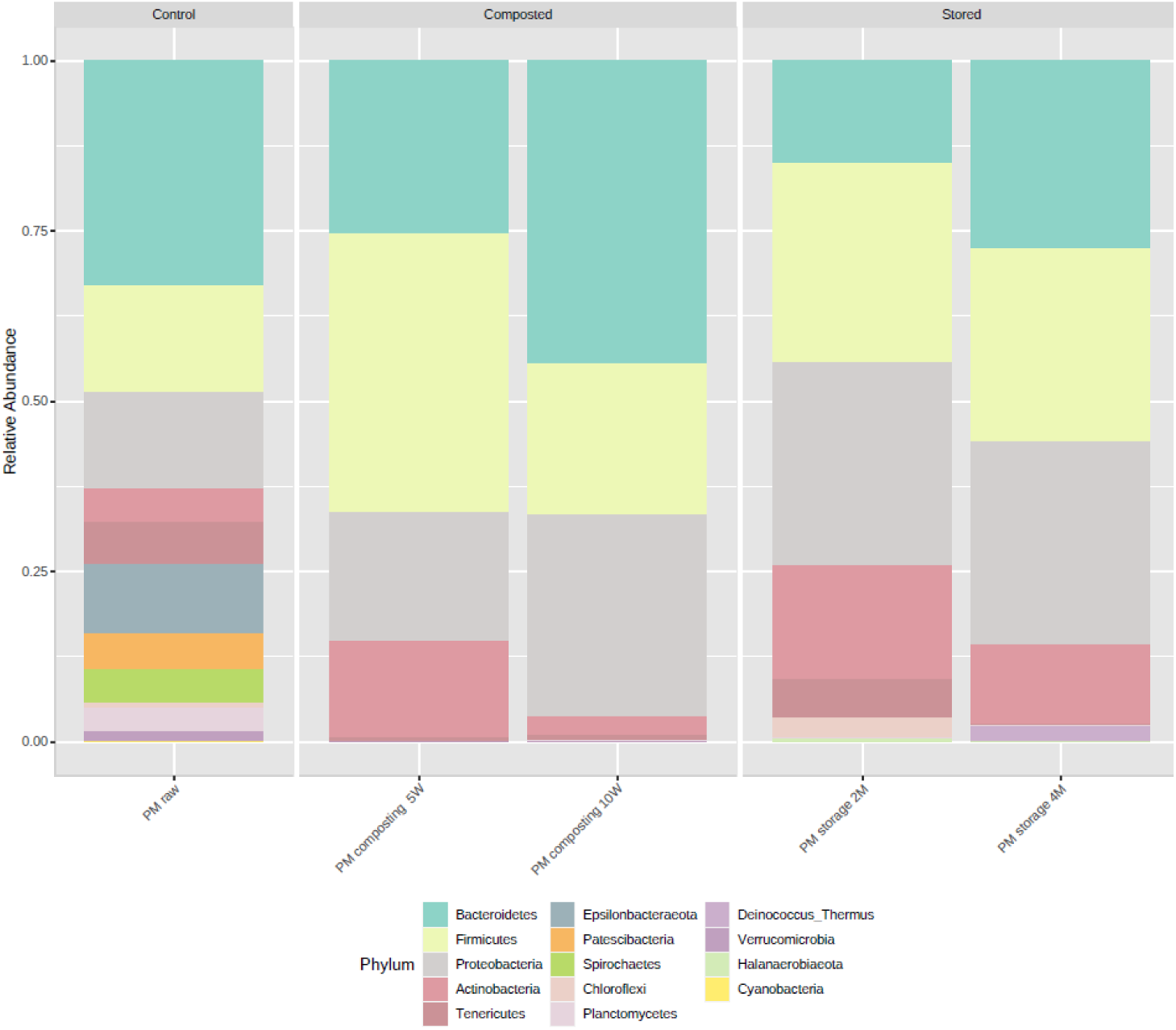
Microbial composition of samples at the phylum level. The samples were divided into three groups – control (PM raw; sample before treatment), composted (PM composting 5W, PM composting 10W; composted samples after five weeks and ten weeks, respectively), and stored (PM stored 2M, PM stored 4M; stored samples after two months and four months, respectively); stacked barplots represents average value for three replicates

At baseline, the most prevalent phylum was *Bacterioidetes* (30%), followed by *Firmicutes* (14%), *Proteobacteria* (13%) and *Actinobacteria* (5%). The following phyla were also abundant: *Epsilonbacteraeota* (9%), *Cloacimonetes* (4%), *Lentisphaerae* (2%), but their abundance differed slightly between samples. The bacterial composition changed after treatment; however, the nature of the changes differed between composting and storage.

The relative abundance of *Bacterioidetes* decreased after five weeks (18.8%) and then increased after ten weeks (43.6%) during composting; however, this value decreased after two months (14.7%) then increased at four months (51.3%) during storage. The relative abundance of *Firmicutes* increased after five weeks (30.2%) and decreased after ten weeks (21.7%) during composting but continuously increased during the storage (28.5% and 52.8, after two and four months, respectively). The relative abundance of *Proteobacteria* increased during both composting (13.8% after five weeks and 28.8% after ten weeks) and storage (28.2% after two months and 55.5% after four months). Finally, the prevalence of *Actinobacteria* initially increased (10.6%) but later decreased at the latest stages (3%) during composting; however, its level increased continuously from 16.4% after two months and 21.8% after four months during storage.

### 3.2. Microbial community diversity

The richness and diversity of the microbial communities were analysed using the Shannon and Chao1 indexes (Fig. 2A and Fig. 2B). The Shannon index showed lower pig manure values during storage and composting than the untreated control samples, but only as a trend. Chao1 index also showed lower pig manure values during storage and composting than the untreated control samples at the trend level. A principal coordinate analysis (PCoA) was used to determine the differences between all samples based on the Bray-Curtis dissimilarity index; the PCoA plot did not show any differences between samples depending on management strategies (untreated manure included).

**Figure 2:**
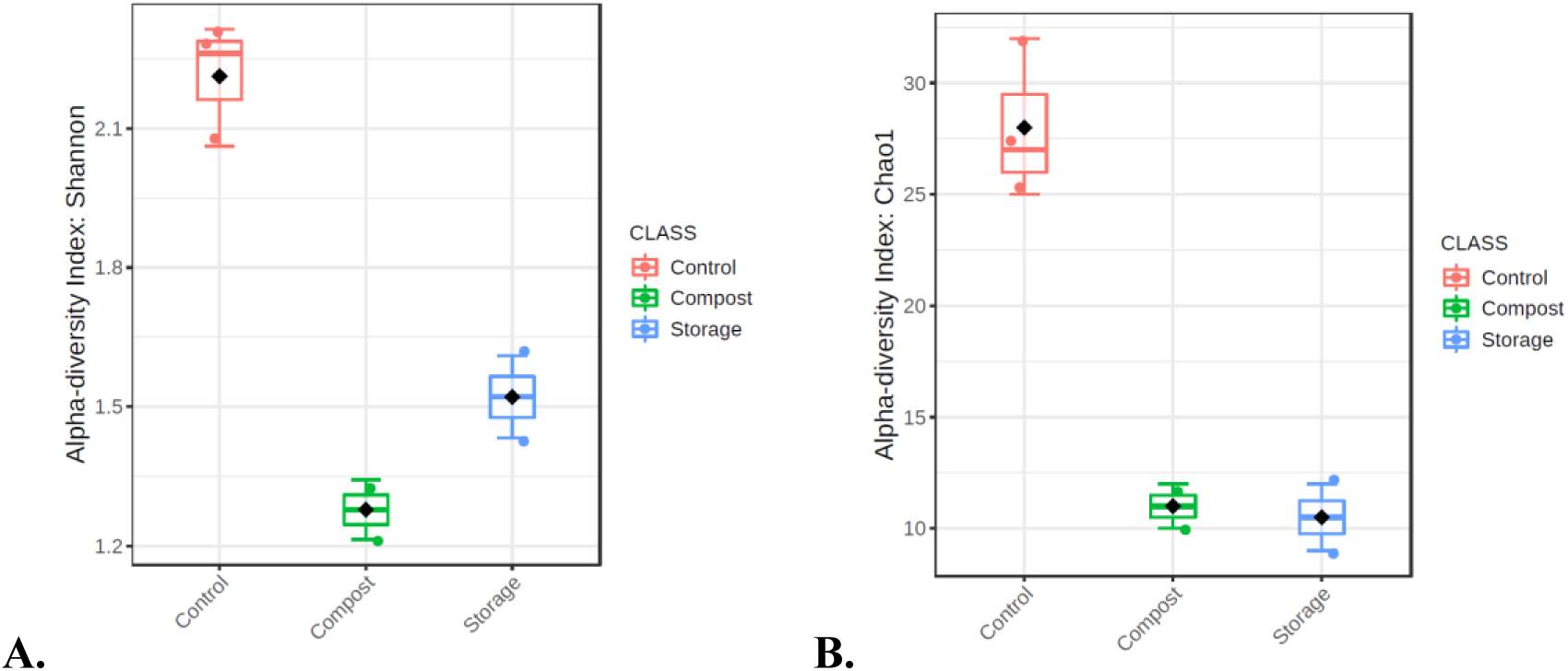
**A.** Shannon indexes of microbial community diversity (p-value 0.068661 [Kruskal-Wallis]) **B.** Chao 1 indexes of microbial community richness (p-value 0.098452 [Kruskal-Wallis]); the samples were divided into three groups – control (PM raw 1, PM raw 2, PM raw 3; samples before treatment), composted (PM composting 5W, PM composting 10W; composted samples after five and ten weeks, respectively), and stored (PM stored 2M, PM stored 4M; stored samples after two and four months, respectively)

### 3.3. Diversity and abundance of ARGs

The relative abundances of the analysed gene classes and MGEs varied during composting and storage (Fig. 3). In both cases, a higher total abundance of detected genes was observed in treated compared to untreated manure; however, the distribution of individual ARGs and ARG groups was found to differ depending on treatment type. The changes in the presence of individual genes are visualized by a heatmap (Fig. 4). In raw manure, the highest number of detected genes were associated with tetracycline resistance, followed by aminoglycoside, MLSB, and sulfonamide resistance. The lowest abundances were detected for phenicol-, vancomycin-, and trimethoprim-resistance genes (Fig. 3).

**Figure 3:**
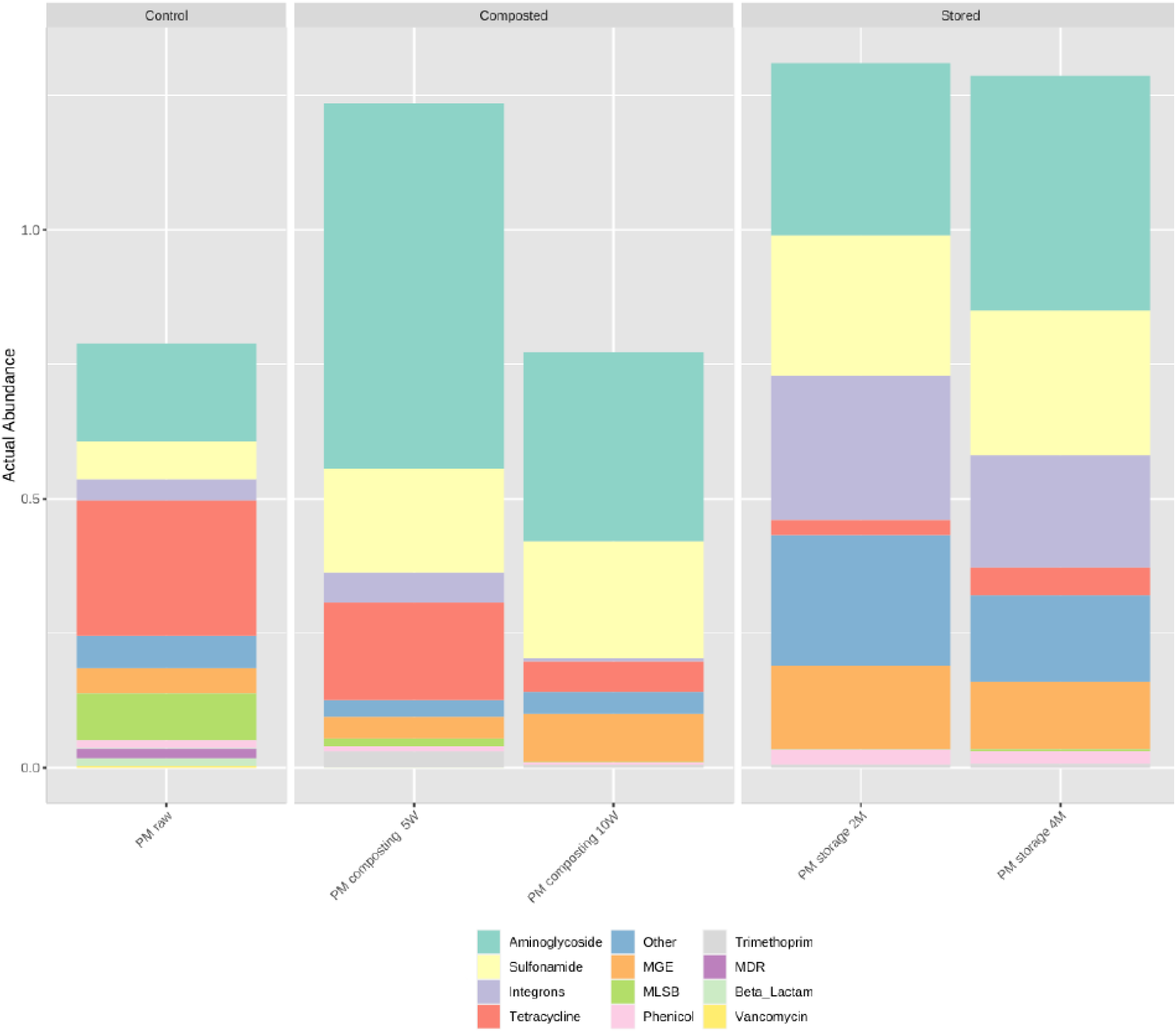
Actual abundance of detected antimicrobial resistance genes and mobile genetic element gene classes detected during qPCR analysis. The samples were divided into three groups – control (PM raw; sample before treatment), composted (PM composting 5W, PM composting 10W; composted samples after five and ten weeks, respectively), and stored (PM stored 2M, PM stored 4M; stored samples after two and four months, respectively); stacked barplots represents the average value for three replicates

**Figure 4:**
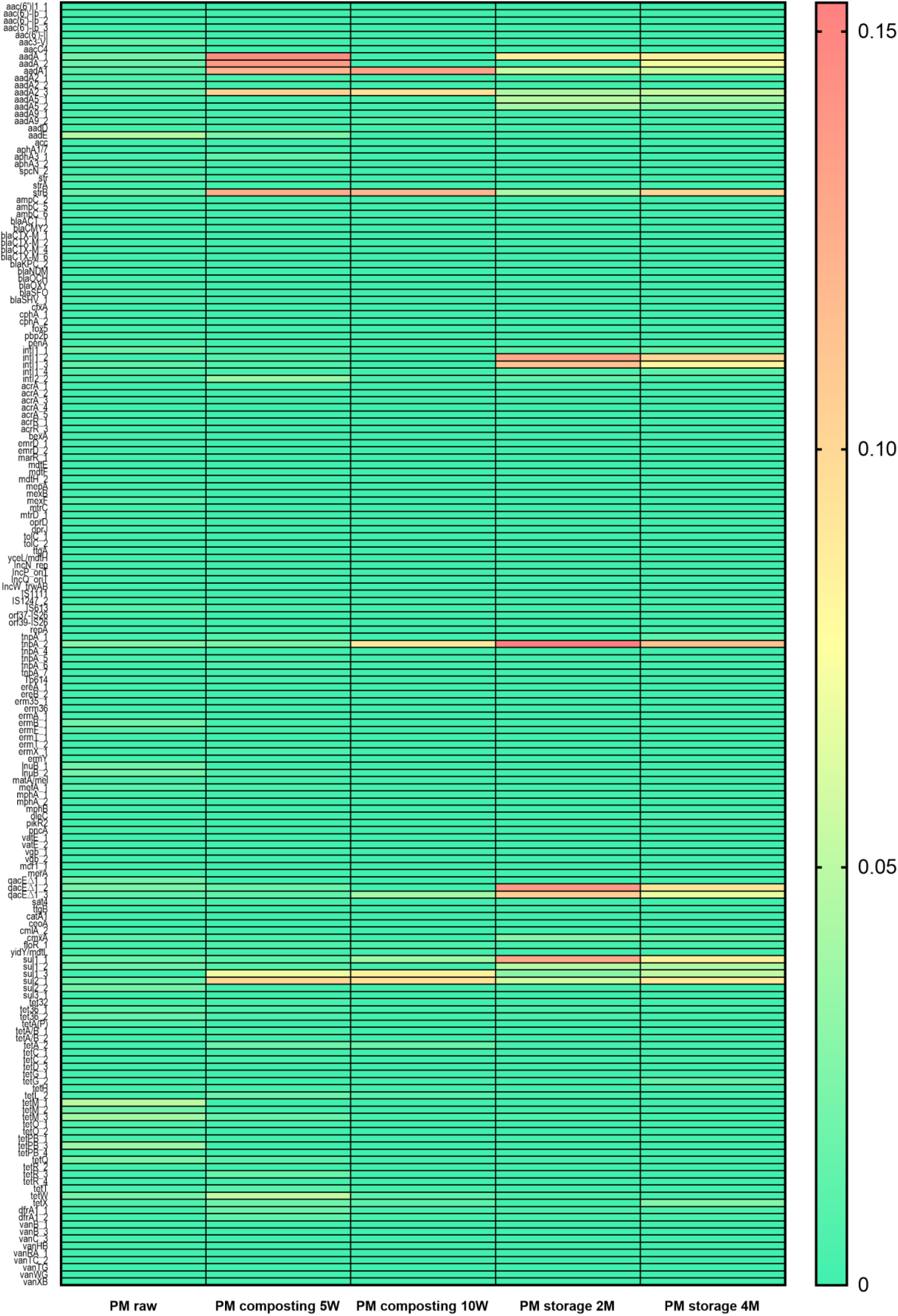
Changes in detected ARG and MGE relative abundances depending on manure treatment strategies compared with control. The samples were divided into three groups – control (PM raw, sample before treatment), composted (PM composting 5W, PM composting 10W; composted samples after five and ten weeks, respectively), and stored (PM stored 2M, PM stored 4M; stored samples after two and four months, respectively); heatmap represents average value for three replicates

Composting causes a temporary increase in the total amount of detected genes to 1.23 gene copies/16S rRNA gene copies after five weeks, followed by a decrease to 0.77 gene copies/16S rRNA gene copies after ten weeks. The composting process caused a decline in the total number of tetracycline-resistance genes; it also resulted in a decrease in the numbers of MLSB-, beta-lactam-, phenicol-, and vancomycin-resistance genes, as well as genes known to offer resistance against other antimicrobials. Composting also caused a reduction in the relative amounts of genes responsible for multi-drug resistance (MDR), as well as in the relative abundance of genes classified as integrons. However, the process resulted in an increase in the total amounts of aminoglycoside-resistance genes and sulfonamide-resistance genes, as well as the amount of MGEs.

In contrast, during storage, the relative abundance of genes increased to 1.31 gene copies/16S rRNA gene copies after two months, and then decreased after four months (1.29 gene copies/16S rRNA gene copies). The storage process caused a decline in tetracycline-, MLSB-, beta-lactam-, and vancomycin-resistance genes, and many aminoglycoside-, sulfonamide-, phenicol-, and trimethoprim-resistance genes were detected after four months of treatment. Storage also caused an increase in the number of resistance genes against other antimicrobials, integrons and MGEs. For both processes, the number of detected genes remained elevated compared to the initial level of 0.79/16S rRNA gene copies. However, in the case of composting, all genes reached their lowest levels after week 10.

While the relative abundance of ARGs was found to generally increase during both treatment strategies this tendency varied depending on the gene class (Tab. S2). The relative abundance of beta-lactam-, tetracycline-, vancomycin-resistance genes, genes conferring the MLSB resistance phenotype, and genes coding the MDR phenotype were reduced at the end of composting and at the end of storage. Interestingly, while the relative abundance of integrons decreased during composting, their copy number increased during storage. A similar situation was observed with phenicol-resistance genes: their abundance decreased after composting but increased after storage for four months. In addition, resistance genes classified as *other* decreased after composting and increased after storage. The number of copies of the remaining analysed ARGs groups increased during both manure management strategies, but the size degree of the change varied according to the process.

The PCoA the relative abundances of ARGs and MGEs in all sample groups, calculated based on the Bray-Curtis distance method, revealed significant visual separation between the initial (untreated) and treated samples with regard to the relative abundances of resistant gene classes (p<0.05) (Fig. 5).

**Figure 5:**
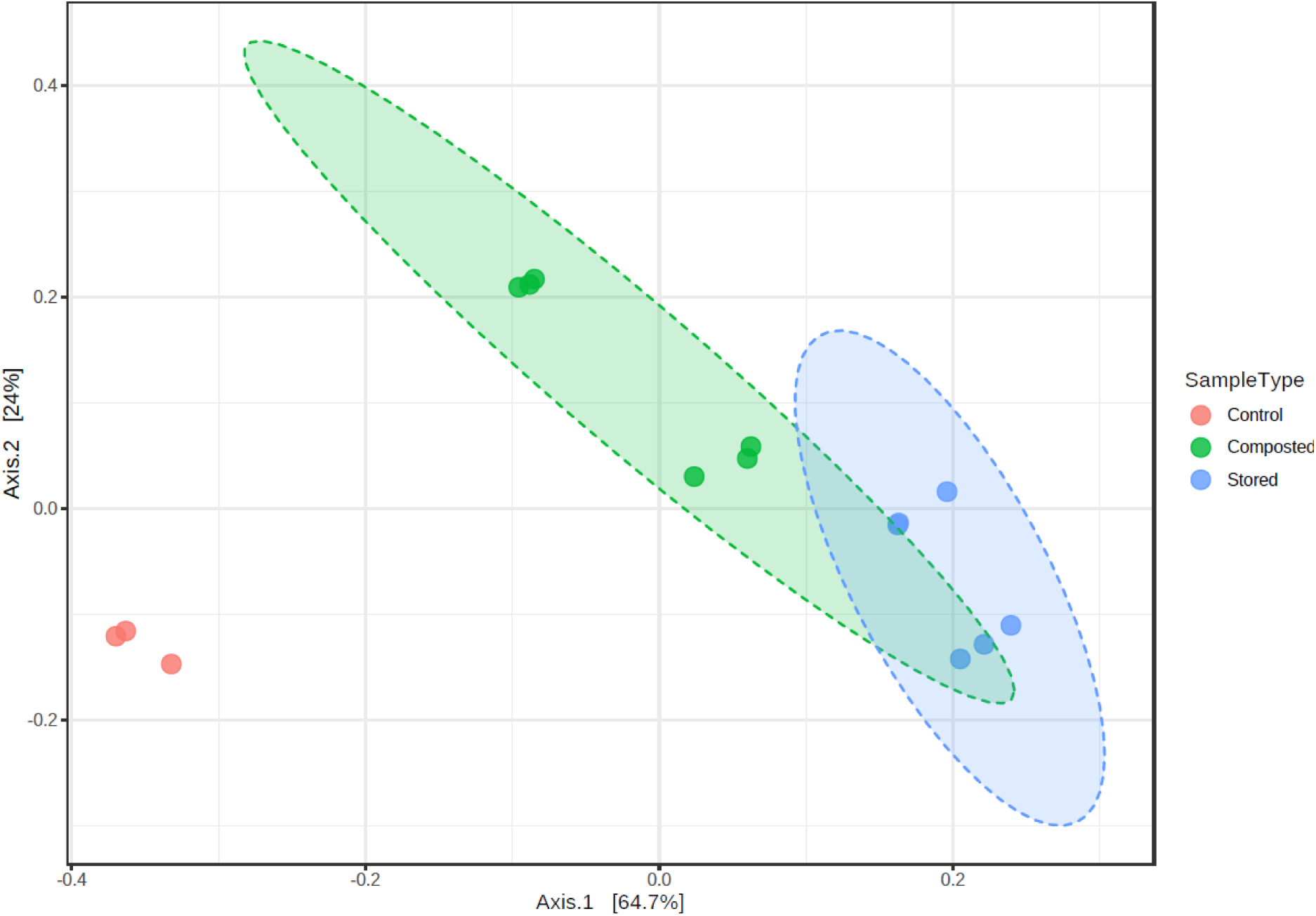
Principal coordinate analysis (PCoA) of ARG and MGE relative abundances in all sample groups based on the Bray-Curtis distance method. The ANOSIM statistical method was used. [ANOSIM] R: 0.98401; p-value <0.001. The samples were divided into three groups – control (PM raw 1, PM raw 2; samples before treatment), composted (PM composting 5W, PM composting 10W; composted samples after five and ten weeks, respectively), and stored (PM stored 2M, PM stored 4M; stored samples after two and four months, respectively)

The most prevalent gene in raw pig manure was *aadE*, followed by *tetM_1.* Both genes were effectively reduced during both composting and storage. Of the ten most common genes in raw pig manure (*aadE*, *tetM_1*, *tetM_3*, *tetPB_3*, *tnpA_2*, *qacEΔ1_2*, *tetQ*, *sul2_2*, *tetW*, *tetM_2* – ranked from high to low), two increased after treatment: *tnpA_2*, and *qacEΔ1_2.* The former belongs to the IS21 group of transposases and was assigned during this study to the MGE group, while the latter codes efflux pumps, which is one of the mechanisms responsible for MDR. Both composting and storage caused a reduction in the abundance of 80% of the genes detected in raw manure; of these, one is an aminoglycoside-resistance gene, six are tetracycline-resistance genes, one was a mobile genetic element, one an MDR mechanism, and one a sulfonamide resistance gene. The core resistome of the analysed samples is presented in fig. 6.

**Figure 6:**
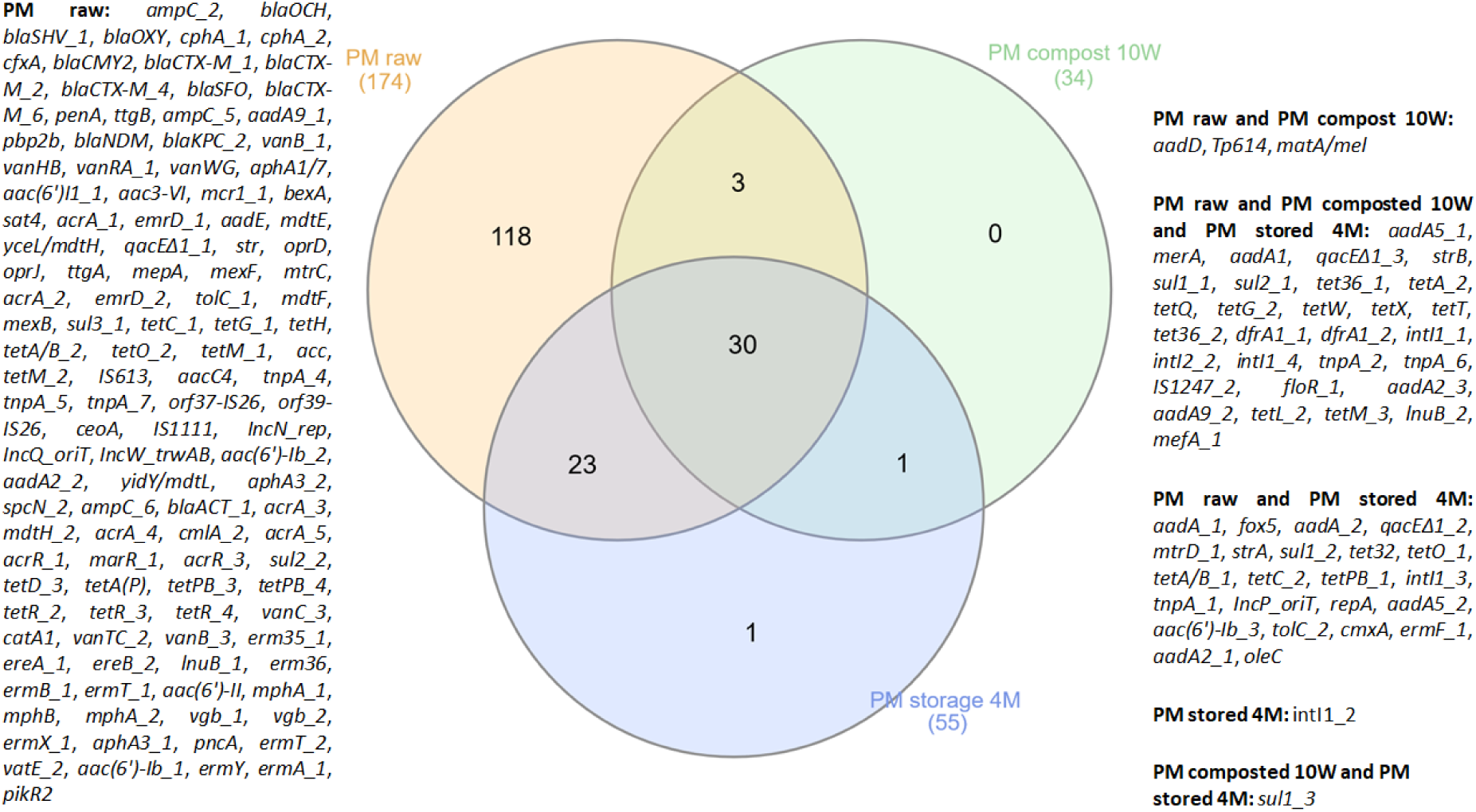
Core resistome in pig manure samples. The Venn diagram, presenting the number of detected genes across all samples and their intersection, was created by InteraciVenn (Heberle et al., 2015). The samples were divided into three groups: PM raw - untreated pig manure, PM compost 10W - pig manure after 10 weeks of composting, PM storage 4M - pig manure after four months of storage

The applied manure management strategies were found to only partially reduce ARG diversity (Tab. S3). The number of integrons increased during four-month storage, but decreased during composting. The number of trimethoprim-resistance genes was not reduced during either procedure. The numbers of aminoglycoside-, beta-lactam-, phenicol-, sulfonamide-, tetracycline-, vancomycin-resistance genes, genes coding the MLSB resistance phenotype, genes conferring the MDR phenotype, MGEs, resistance genes against other antimicrobials were decreased.

### 3.4. Correlation analysis of ARGs, MGEs, and microbial communities across the manure treatment processes in pig manure

A co-occurrence network was constructed to determine the relationship between 157 ARGs and 21 MGEs (integrons included) during composting. This network (Fig. 7) contains, altogether, 179 nodes, and was created based on 959 strong (|r|>0,9) and significant correlations (p≤0.05) with 942 positive correlations and 17 negative ones. The MGEs showed more connections than ARGs, indicating a more important role in network formulation. Among the MGEs and integrons group, *orf39-IS26*, *IncN_rep*, *IncQ_oriT*, *orf37-IS26*, *IS1111*, *tnpA_4*, *IncW_trwAB* showed the highest number of correlations with ARGs. Among the analyzed MGEs and integrons, only *IncP_oriT*, *IS613*, *intI1_4*, *intI1_1*, *repA*, *tnpA_1*, *Tp614*, *IS1247_2*, *tnpA_2* demonstrated positive and negative interactions: the rest of genes assigned to this group only demonstrated positive correlations (*orf39-IS26*, *IncN_rep*, *IncQ_oriT*, *orf37-IS26*, *IS1111*, *tnpA_4*, *IncW_trwAB*, *tnpA_5*, *intI1_3*, *IncP_oriT*, *IS613*, *intI1_4*, *intI1_1*, *repA*, *tnpA_1*, *Tp614*, *IS1247_2*, *tnpA_2*, *intI2_2*, *tnpA_7*, *intI1_2*). None of the analysed MGEs or integrons displayed only negative correlations.

**Figure 7:**
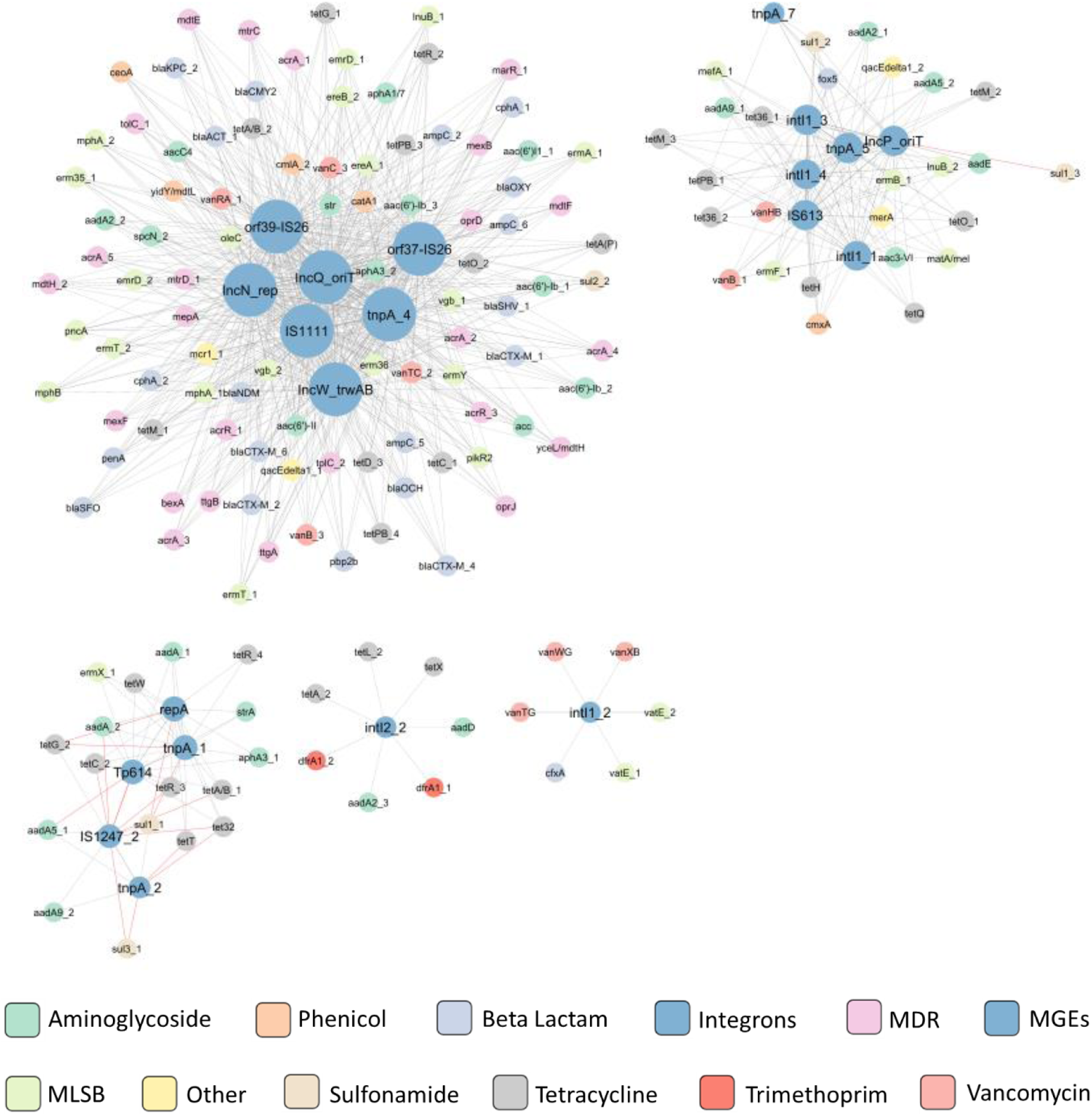
ARG and MGE interaction network in compost. The network is presented as an ’organic layout’. A strong and significant correlation is shown, based on Spearman’s rank correlation, where |r|>0.9 and p<0.05. The size of the nodes represents the degree of interaction. The gray and orange edges show positive and negative correlations between ARGs and MGEs, respectively. The color of the nodes indicated the ARG type according to the legend

The relationship between 155 ARGs and 19 MGEs (together with integrons) during storage was determined by constructing a co-occurrence network. The analysis comprised 155 ARGs and 19 MGEs assessed across the storage process. This network (Fig. 8) contained a total of 152 nodes, and it was created based on 1224 strong (|r|>0,9) and significant correlations (p≤0.05) with 1211 positive correlations and 13 negative ones. The MGEs showed more connections than ARGs, indicating they play an important role in network formulation. Among the MGE and integron group, *orf39-IS26*, *IncN_rep, IncQ_oriT*, *orf37-IS26, IS1111*, *tnpA_4*, *IncW_trwAB*, *tnpA_5 tnpA_7*, *intI1_3*, *IS613*, *intI1_4*, intI1_1, *repA*, *tnpA_1*, *IS1247_2*, *tnpA_2*, *intI2_2*, showed the highest number of correlations with ARGs. Among this group, only *intI1_3*, *IS613*, *intI1_4*, *intI1_1*, *repA*, *tnpA_1*, *IS1247_2*, *tnpA_2*, *intI2_2*, *tnpA_7*, *intI1_2* possessed positive and negative interactions: *orf39-IS26*, *IncN_rep*, *IncQ_oriT*, *orf37-IS26*, *IS1111*, *tnpA_4*, *IncW_trwAB*, *tnpA_5*, *intI1_3*, *IS613*, *intI1_4*, *intI1_1*, *repA*, *tnpA_1*, *IS1247_2*, *tnpA_2*, *intI2_2*, *tnpA_7* only demonstrated positive correlations. Like composting, none of the analysed MGEs or integrons showed only negative correlations.

**Figure 8:**
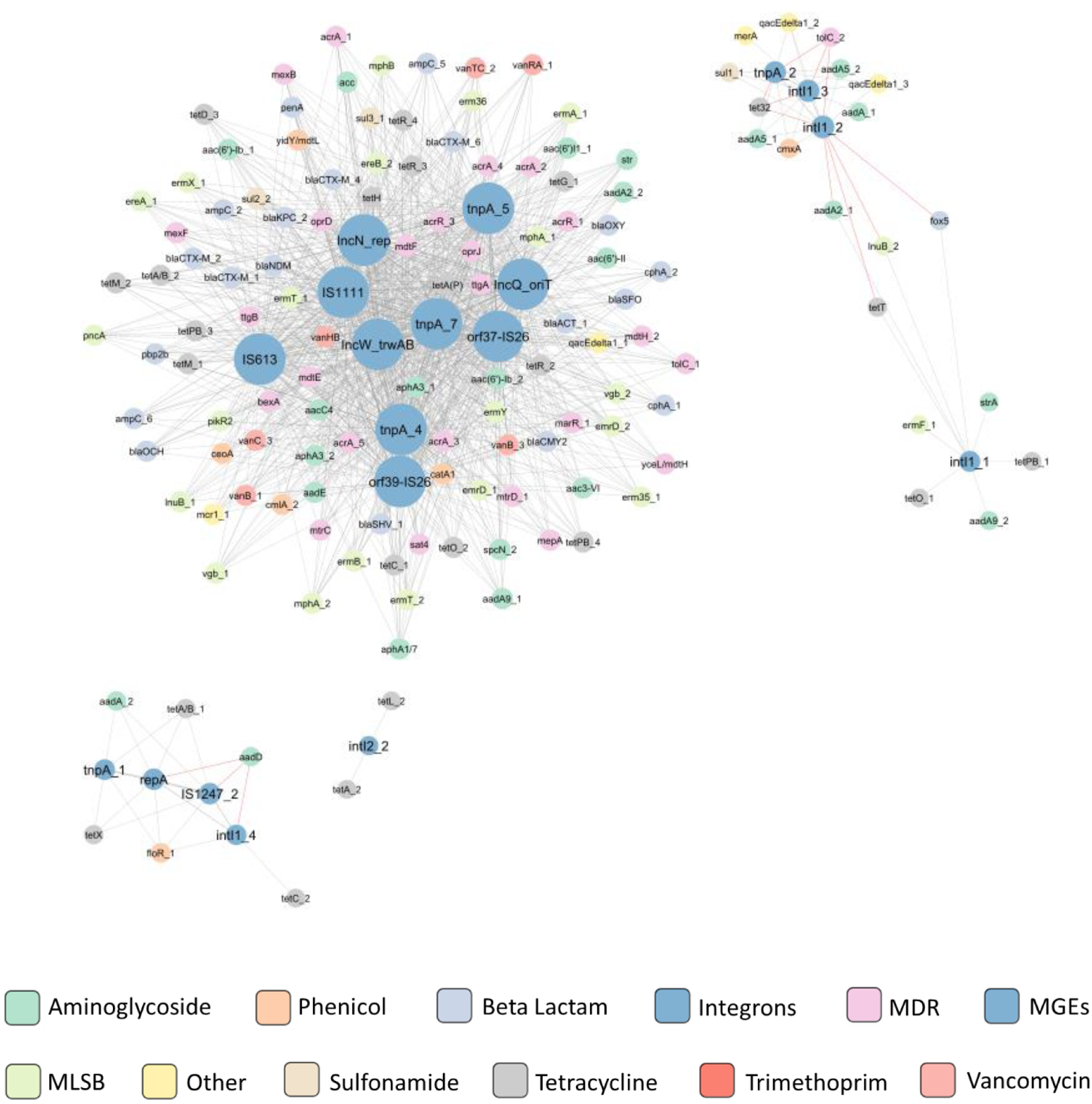
ARG and MGE interaction network in compost. The network is presented as an ’organic layout’. A strong and significant correlation is shown based on Spearman’s rank correlation, where |r|>0.9 and p<0.05. The size of the nodes represents the degree of interaction. The gray and blue edges show positive and negative correlations between ARGs and MGEs, respectively. The color of the nodes indicates the ARG type according to the legend

### 3.5. Correlation between detected genes in HT-qPCR and microbial taxa

The interactions between 26 phyla and 12 gene classes (10 ARGs classes, integrons, and MGEs) identified during the composting procedure were investigated. The resulting co-occurrence network (fig. 9) contained 37 nodes, and it was created based on 79 strong (|r|>0,85) and significant (p≤0.05) correlations with 54 positive and 25 negative correlations. As shown in the network, the gene classes presented more interactions than the microbial phyla, indicating that these genes play a crucial role in the co-occurrence network construction. While the genes coding the MDR mechanism demonstrated the highest interaction with microbial phyla, genes coding the resistance to aminoglycosides, trimethoprim, and genes assigned as *Other* class also exhibited high levels of interaction. The lowest level of connections were observed for integrons and MGEs. The MDR genes only demonstrated positive correlations with ten microbial phyla: *Artibacteria*, *Chlamydiae*, *Cloacimonetes*, *Elusimicrobia*, *Kiritimatiellaeota*, *Lentisphaerae*, *Marinimicrobia*, *Synergistetes*, *Thermotogae*, and *Fibrobacteres*. Other ARG groups show both positive and negative correlations with microbial phyla. The co-occurrence networks created by MGE and integrons, and by genes conferring resistance to *inter alia* aminoglycosides and trimethoprim, were separated from those created by genes conferring the MDR type and genes conferring resistance to beta-lactam, tetracycline, sulfonamide, phenicol, MLSB and vancomycin.

**Figure 9:**
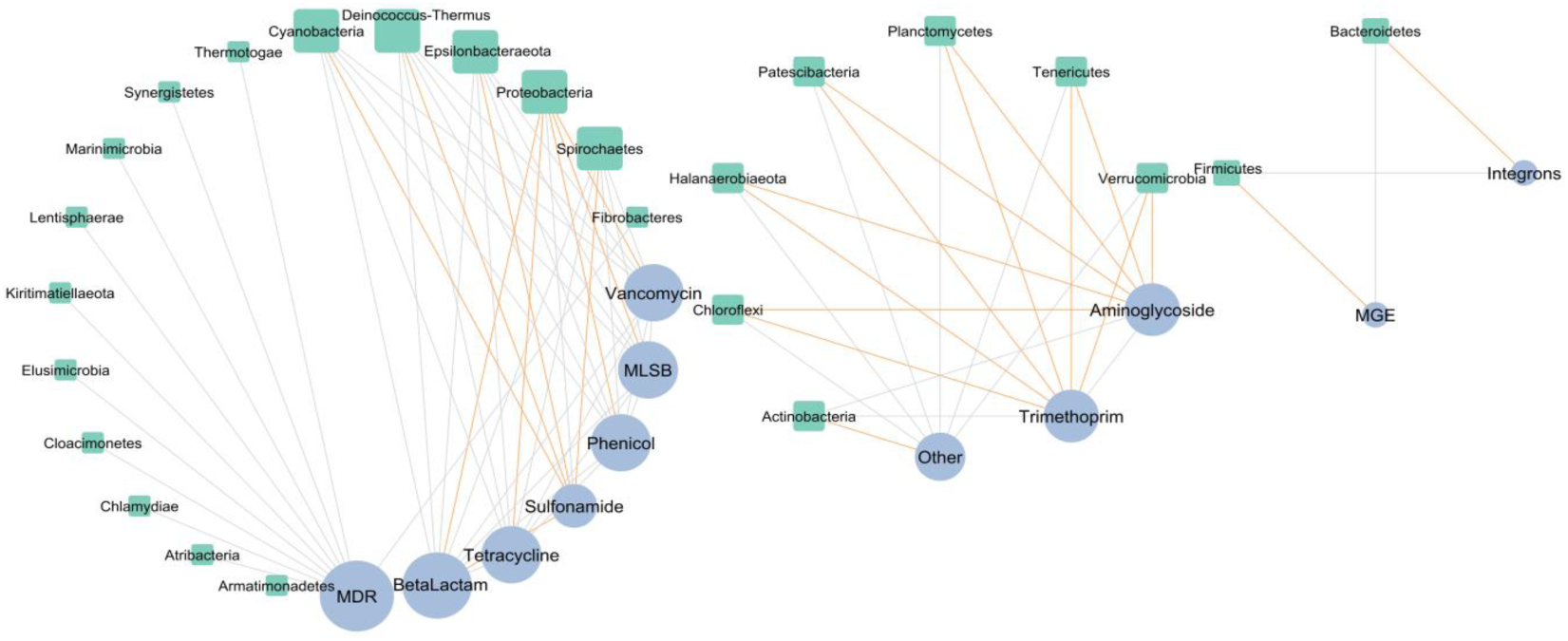
Interaction network analysis of microbial taxa and gene classes during composting, presented as a ’circular layout’ based on the co-occurrence of ARG groups and microbial phyla. A connection is significantly correlated with Spearman’s |r|>0.85 and p<0.05. The gray and orange edges represent the positive and negative correlations between ARG groups and phyla, respectively. The size of the nodes shows the degree of interaction. Green nodes represent microbial phyla, and blue nodes represent groups of ARGs

Among the microbial taxa, *Cyanobacteria*, *Deinococcus-Thermus*, *Epsilonbacteriaeota*, *Proteobacteria* and *Spirochaetes* demonstrated the most interactions with ARG groups (negative and positive); however, they appear to interact only with genes conferring MDR and MLSB resistance, and resistance to beta-lactam, tetracycline, sulfonamide, phenicol and vancomycin. *Firmicutes* and *Bacteroidetes* were clustered with MGE and integrons in a separate network. Similarly, *Actinobacteria*, *Chloroflexi*, *Halanaerobiaeota*, *Patescibacteria*, *Planctomycetes*, *Tenericutes*, and *Verrucomicrobia* were clustered separately from other phyla and interacted with genes classified as *other*, as well as those providing trimethoprim and aminoglycoside resistance.

The interactions between 26 phyla and 12 gene classes (10 ARG classes, integrons and MGEs) identified during pig manure storage were investigated. The constructed co-occurrence network (fig. 10) contained 34 nodes, and was created based on 58 strong (|r|>0.85) and significant (p≤0.05) correlations: 40 positive and 18 negative correlations. As shown in the network, the gene classes presented more interactions than the microbial phyla, indicating the crucial role played by these genes in the construction of the co-occurrence network. Of these genes, the vancomycin-resistance group demonstrated the most interaction, creating a separate cluster with the following phyla: *Armatimonadetes*, *Atribacteria*, *Chlamydiae*, *Cloacimonetes*, *Cyanobacteria*, *Elusimicrobia*, *Fibrobacteres*, *Kiritimatiellaeota*, *Lentisphaerae*, *Marinimicrobia*, *Synergistetes*, *Thermotogae*, and *Verrucomicrobia*. Vancomycin only demonstrated positive interactions with these phyla. Similarly, only positive interactions were observed with genes conferring MLSB resistance mechanisms and tetracycline-resistant genes, which belong to separate clusters. Other gene classes demonstrated positive and negative interactions. Only the sulfonamide-resistance genes showed no interactions with these bacterial phyla.

**Figure 10:**
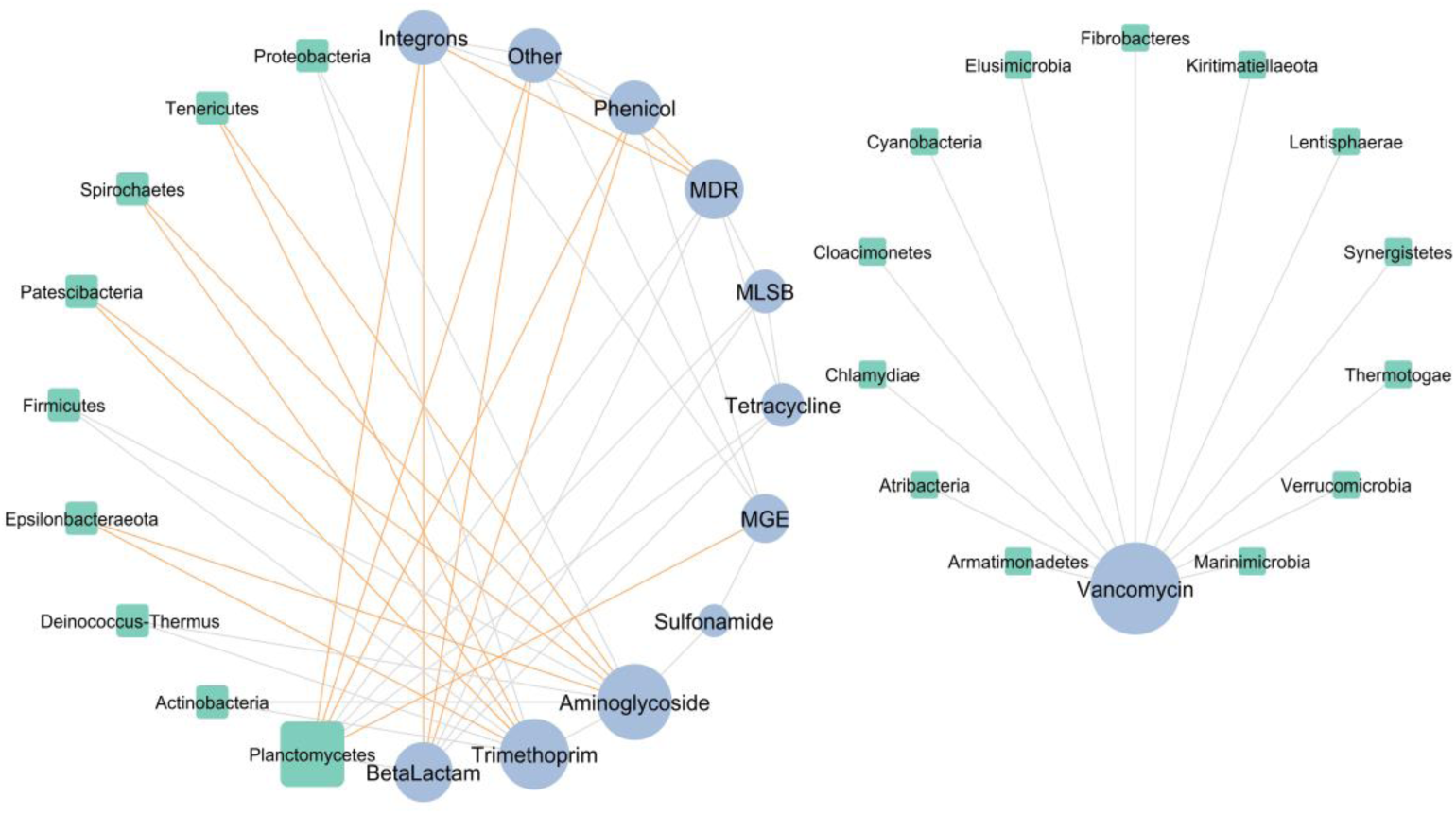
Interaction network analysis of microbial taxa and gene classes during storage, presented as a ’circular layout’ based on the co-occurrence of ARG groups and microbial phylum. A connection is significantly correlated when Spearman’s |r|>0.85 and p<0.05. The gray and orange edges represent the positive and negative correlations between ARG groups and phyla, respectively. The size of the nodes shows the degree of interaction. Green nodes represent microbial phyla, and blue nodes represent groups of ARGs

Apart from *Planctomycetes*, all bacterial phyla demonstrated either negative or positive correlations with the studied gene classes. Of the phyla, *Planctomycetes* demonstrated the most correlations, i.e. three positives (MDR, MLSB, and tetracycline-resistance genes) and four negatives (MGE, integrons, genes classified as *other-*, and phenicol-resistance genes).

## 4. Discussion

### 4.1. Raw pig manure as a potential source of contamination in the natural environment

Livestock manure has been recognized as an essential hot spot for the dissemination of antimicrobial resistance into the natural environment. This study examines the extent of the ARG reservoir in medium-scale animal production facilities. The most common antibiotics used in pig production are tetracyclines, macrolides and sulfonamides (R.-M. Zhang et al., 2021). This was in line with our findings, indicating that resistance to these antibiotics predominated among the detected ARGs. These findings are consistent with previous studies (Xu et al., 2020; Zhu et al., 2013), which identified macrolide-, cephalosporin-, aminoglycoside-, and tetracycline-resistance genes as the most prevalent types.

The diverse set of ARGs determined during the present study (Fig. 6) potentially confer resistance to all major classes of antibiotics, including antibiotics that are critically important for human medicine (WHO Advisory Group on Integrated Surveillance of Antimicrobial Resistance and World Health Organization, 2017), such as macrolides (*erm* and *aph* genes), beta-lactams (bla_SHV_, *bla_OXY_*, *bla_CTX-M_*), aminoglycosides (*aad*), sulfonamides (*sul*), and tetracycline (*tet*). Although a few vancomycin-resistance genes were identified (*vanB*, *vanC*) in the raw manure samples, the genes were only detected at low levels and only a partial gene presence was observed (*vanHB* or *vanXB*), assuming that resistance requires the presence of multigene *van* operons (*vanH-vanA-vanX*) (D’Costa et al., 2006), it is not anticipated that the bacteria demonstrate significant phenotypic resistance to vancomycin. However, it is possible that enrichment for van operons can occur under favourable conditions, as indicated by the detection of individual vancomycin-resistance genes. Genes potentially conferring resistance to aminoglycosides, tetracyclines, sulfonamide, phenicols, beta-lactams, macrolides, or quaternary ammonium compounds were detected in all raw manure samples.

Together with sulfonamides, tetracycline is one of the most commonly-applied antibiotics to livestock. They offer a broad spectrum of activity, low toxicity and low price; however, after administration, most tetracyclines are excreted in still active form. In our study, the most prevalent in pig manure were *tetM, tetPB, tetQ, tetW,* and *tet36*, with *tetO, tetT, tetD, tetR* and *tet32* being found in lower amounts. Zhu et al. (Zhu et al., 2013) found *tetQ, tetW, tetX, tet(32), tetO, tetM, tetL*, and *tetG* to be the most abundant tet genes in manure, while Lim et al. (Lim et al., 2020) found *tetQ, tet32, tet44,* and *tetW* to be the most prevalent and abundant genes; the *tet32* family was also found to be of the most prevalent among swine in a previous study (Li et al., 2015). Ghosh and LaPara (Ghosh and LaPara, 2007) found the most common to be *tetL, tetA, tetM*, and *tetG.* Interestingly, *tet32, tetO, tetQ,* and *tetW* were identified as the most prevalent genes in humans in China, Denmark, and Spain (Forslund et al., 2013; Hu et al., 2013).

Beta lactams are also commonly administered to pigs. Among the known beta lactam-resistance genes, *bla_SHV_*, *penA*, *fox5*, *pbp2*, *bla_OXY_*, *bla_CTX-M_* were found to be the most prevalent in raw pig manure in the present study. bla_SHV_, *bla_OXY_*, *bla_CTX-M_* are the most ESBL-coding genes, and their presence has been confirmed in *Enterobacteriaceae* (e.g., *E. coli* or *K.* pneumoniae) isolated from many farm animals (Ejaz et al., 2021; Gao et al., 2015; García-Cobos et al., 2015); however, previous studies have often identified *bla*_TEM_, which we do not observed in the present study. Gao et al. (Gao et al., 2015) report the presence of *E. coli* strains harbouring both *bla_CTX-M_* and *bla*_TEM_ in pig manure. One beta-lactamase originally found in *K. pneumoniae* is FOX-5 (*fox5*), known to be responsible primarily for resistance to cephamycin and cephalosporin.

Our present findings also identified genes of special concern, such as *bla_NDM_* (New Delhi metallo-β-lactamase1) and *bla_KPC_* (isolates that produce *K. pneumoniae* carbapenemase), albeit at relatively low intensity. The co-occurrence analysis indicated a positive correlation with almost all identified MGEs, implying high mobility and a high risk of potential HGT. Infections caused by pathogens harbouring these genes are especially threatening due to limited treatment options (Zalewska et al., 2021). Although these genes have been reported worldwide in healthcare facilities, little is known of their prevalence in a swine production environment (Taggar et al., 2020).

The most abundant macrolide-resistance genes identified during our study were *lnuB* encoding lincosamide nucleotidyltransferase, responsible for clindamycin resistance in patients hospitalized with *Streptococcus* spp. *or Staphylococcus* spp. (Almuzara et al., 2013)*, ermB* conferring erythromycin resistance in a variety of gram-positive bacteria (e.g., *L. monocytogenes*) (Gupta et al., 2003), *mefA* responsible for macrolide resistance in pneumococci in hospitalized patients in Europe (Ardanuy et al., 2005), *ermF*, conferring MLSB resistance in *Bacterioidetes*, often located on MGEs (Chung et al., 1999), and *mphA* encoding resistance to azithromycin in enterotoxigenic *E. coli* (Xiang et al., 2020). Macrolide-resistant bacteria, carrying erythromycin ribosome methylation (*erm*) genes and macrolide efflux (*mef*) genes, are also excreted in faeces. Moreover, many *erm* genes have been found in swine waste lagoons: *ermA*, *ermB*, *ermC*, *ermF*, *ermG*, *ermT*, *ermQ,* and *ermX* have been identified, with *ermB* being the most prevalent, followed by *ermF* (Garder et al., 2014; Luby et al., 2016).

The most abundant aminoglycoside-resistance genes identified during this study were *aadE* (confers streptomycin resistance) (Lysnyansky and Borovok, n.d.), *aadA* (Streptomycin/Spectinomycin Adenylyl transferase) (Clark et al., 1999), *aadA2* (streptomycin resistance) (Briggs and Fratamico, 1999)*, strB* (streptomycin resistance) (Sundin and Bender, 1996). Moreover, highly-transposable *aphA3* was also identified, known to be responsible for kanamycin resistance (Lysnyansky and Borovok, n.d.). Our results confirm previous findings by other research teams, e.g., Xiong et al. (Xiong et al., 2019), with the aminoglycoside resistance genes *aadA* and *aph1* being present in all analysed swine manure samples, and *aadA* being one of the most prevalent genes in all manure samples.

Heavy metals are common in the natural environment and may prevent ARG removal, even under favourable conditions. In agricultural practice, heavy metals including zinc, copper, lead, cadmium, chrome, and nickel have been commonly found in animal faeces. Small doses of zinc and copper are added as mineral supplementation to animal food to enhance growth (Poole, 2017), and lead, cadmium, chrome, and nickel are introduced to manure from corroded machinery (Zhao et al., 2018). Even if heavy metal ions are present in manure in an acceptable range covered by regional regulations, such sublethal doses facilitate the maintenance of ARGs (Hölzel et al., 2012) and support the conjugative transfer of ARGs (Zhang et al., 2018). Our present findings indicate the presence of the *merA* gene coding mercuric reductase, responsible for mercury resistance; it may serve as a marker of the possible co-occurrence of ARGs and heavy metal resistance genes. However, Zhang et al. (Zhang et al., 2018) report that environmental factors such as temperature, pH, ammonia content, and moisture have a greater influence on ARG profile dynamics than the presence of heavy metals or integron, and surprisingly, that the physicochemical properties of compost have a greater impact than the antibiotics; however, the researchers used tylosin during the study, which rapidly degraded during the thermophilic phase of composting.

The differences in ARG profile observed between the treated pig manures may occur due to shifts in the bacterial taxa inhabiting the gastrointestinal tract. These are known to be influenced by various factors, but particularly animal age and type of herd management, i.e. the system of feeding or antibiotic use. Management may be divided into three types: 1) sow (breeding, gestation, farrowing); 2) nursery (21 days, 18 kg feeders); and 3) finisher farms (feeders to 113 kg) (McLaughlin et al., 2009). Animal management, including antibiotics, is carefully employed at each stage to ensure proper growth or treat infections. Brooks et al. (Brooks et al., 2014) found that finisher farm wastewaters harboured the fewest ARGs and represented the least diverse microbial ecology, while sow and nursery farms presented the highest antibiotic resistance. However, Lu and Lu (Lu and Lu, 2019) found animal age to have no observable effect on ARG profile for solid manure.

### 4.2 Manure treatment

The long-term goal of manure management is to remove environmental contaminants: disposing of the high volume of waste, using its value to improve soil fertility, and ensuring biosecurity. Hence, it was necessary to study the physicochemical properties of both treated and untreated manure. The findings regarding the concentration of heavy metals are in line with those obtained by previous research on pig manure, which also proved that the bioavailability of heavy metals decreases during the composting process, with the high temperature during compost being assumed to play a key role (Shehata et al., 2019; Vukobratović et al., 2014). Manure is considered to be a slower fertilizer than slurry, mainly because of the nitrogen available to plants. It has been estimated that 15% of organic nitrogen in the manure is easily available to plants, while up to 35% of organic nitrogen can be mineralized in the first year while up to 50% can be mineralized in the next three to four years (Adesemoye and Kloepper, 2009; Diacono and Montemurro, 2010). Therefore, it is recommended to use pig manure in summer or autumn and slurry in spring before or during ploughing. However, composted manure has an increased nitrogen content, which means that it can also be used in spring.

Composted pig manure has an alkaline pH, which contributes to maintaining or increasing the pH of the soil; this is of great importance for maintaining the proper nitrogen cycle and phosphorus availability, and increasing the absorption of micronutrients by plants (D’Hose et al., 2016; Park et al., 2018). Using composted manure to increase the organic matter of soils has an agronomic justification and is also consistent with the EU’s circular economy action plan (“Circular economy action plan,” n.d.): it improves the physical (soil structure) and physicochemical conditions of the soil by increasing its sorption properties (the role of humus) and organic substance content; this stimulates the soil microbiome, which in turn triggers a further chain of changes beneficial for plants (Veeken et al., n.d.).

To halt the spread of ARGs and potentially pathogenic microorganisms from animal faeces to arable fields and then to crops, various manure pre-treatment strategies have been explored. An important goal in this regard is to identify practices that can decrease their concentrations rather than simply diluting them (Zhu et al., 2013). The abundance of ARGs in livestock animal manure after composting is highly variable. It depends on several essential factors, including the reproduction and death rates of the intestinal microorganisms carrying ARGs, the pressure placed on them by antibiotic residues and heavy metals, favouring the persistence of resistance genes in the bacterial community, and the possibility of horizontal gene transfer between bacteria (Lima et al., 2020).

The composting process seems to be effective in the elimination of antibiotic residues, thus limiting one of the crucial factors in selecting for ARGs; for example, chlortetracycline residue was completely removed from antibiotic-containing swine manure after 21 days, while 17-31% of biologically-active ciprofloxacin remains in compost (Bernal et al., 2009). In addition, beta-lactams have a very sensitive ring-shape structure, which is cleaved by ammonia, phosphate, and hydroxyl ions present in composted manure, resulting in its complete degradation (Kakimoto et al., 2007). Zhang et al. (M. Zhang et al., 2019) found that approximately 64.7% of detectable veterinary antibiotics were efficiently removed from manure during composting. The overall effect of composting on ARG removal rate may be explained by several factors.

When quantifying the antibiotic resistome, it is important to consider that changes in resistance gene abundances may simply be due to changes in underlying microbial structure, since community composition can also influence the composition of resistance genes (Muurinen et al., 2021). In the present study, the bacterial community underwent changes during composting and storage; however, no significant differences were noted in bacterial community richness or differences, with only general trends observed in microbial community diversity and richness. The most prevalent phyla observed across all samples were *Firmicutes*, *Bacteroidetes*, and *Proteobacteria*. These three phyla have been found to be the most abundant in the gut microbiome of pigs (Tang et al., 2020). In both cases, *Bacteroidetes* was observed to dominate in raw manure, with its numbers falling in the middle of the treatment, then rising towards the end; however, while composting resulted in its final level being higher than initial levels, storage only returned this to initial levels. For *Proteobacteria* and *Actinobacteria*, the numbers increased in the middle of both processes; however, the number of *Actinobacteria* declined at the end of composting.

*Proteobacteria*, *Bacteroidetes*, and *Actinobacteria* have all been found in high abundance when composting bulking materials rich in lignose and cellulose (e.g., sawdust, hay, or wheat straw) (Liu et al., 2020). *Firmicutes* has a high tolerance to high temperatures, and hence is more abundant during the thermophilic phase, while *Bacteroidetes* prefers lower temperatures and is more abundant during the cooling phase (Liu et al., 2020). Similar results were obtained by Do et al. (Do et al., 2022), but in liquid pig slurry rather than solid manure; however, other researchers have also confirmed these observations for solid pig manure (Lei et al., 2021; Li et al., 2019; Liu et al., 2021). Although storage is the most commonly applied manure management technology, few studies have examined its effect on the bacterial community. *Firmicutes*, *Actinobacteria*, and *Bacteroidetes* have been found to be the most prevalent bacterial phyla in pig manure during storage under diverse conditions (Lim et al., 2018), which is in agreement with our present findings. Do et al. (Do et al., 2022) also report that these phyla dominate in stored manure at different time points.

The network analysis of bacterial phyla and ARG groups in the composted manure found *Proteobacteria* to co-occur with sulfonamide-resistance genes, *Actinobacteria* with aminoglycoside-resistance genes, *Bacteroidetes* with MGEs, and *Firmicutes* with integrons. Our results partially agree with those of a co-occurrence analysis by Liu et. al. (Liu et al., 2020), who find that all bacterial phyla (*Proteobacteria*, *Bacteroidetes*, Firmicutes, and *Actinobacteria*) that predominated during composting are strongly associated with tetracycline-resistance genes, quinolone-resistance genes, and sulfonamide-resistance genes. In addition, Song et al. (Song et al., 2017) also found these phyla to be associated with macrolide-resistance genes. A similar analysis performed on manure sampled during storage found *Actinobacteria, Firmicutes* and *Proteobacteria* to be associated with that of trimethoprim-resistance genes and aminoglycoside-resistance genes. However, those are only calculated based on Spearman’s correlations and they need to be evaluated by experimentation: other research teams may set different parameters or cut-off points.

Although composting is efficient at antibiotic removal, the situations regarding ARG removal is more complicated. Our study detected an elevated ARG and MGE abundance after all applied manure treatment strategies; however, the smallest increase was observed for samples collected after 10 weeks of composting. Similar increases in resistance gene prevalence has been previously noted by Cao et al. (Cao et al., 2020) and recently by Do et al. (Do et al., 2022). Moreover, in the PCoA plot of ARGs, the cluster assigned to the initial untreated samples was clearly separate from those assigned to treated manure; however, the two clusters belonging to stored and composted samples overlap. Despite this, the efficiency of ARG and MGE removal is not always satisfactory: some studies report an increase in their relative abundances, while others report a decrease (Zhang et al., 2017). A study on swine manure composting yielded the following removal rates for genes divided according to antibiotic groups (Lu and Lu, 2019): beta lactam 36.8-73.7%, FCA 11.8-52.9%, MLSB 0-47.6%, tetracycline 4.5-36.4%, vancomycin 0-71.4%, aminoglycoside 0-26.9% and sulfonamide 0%. In our present study, composting resulted in reductions for beta-lactam- (100%), vancomycin- (100%), MLSB- (86.36%), phenicol- (83.33%), aminoglycoside-(79.31%), other (71.43%), tetracycline- (65.52%), and sulfonamide-resistance genes (40%), with no reduction observed for trimethoprim-resistance genes. These results only partially agree with ours. Although we found an increase in MGE abundance, as reported previously (Do et al., 2022), this increase was smaller after composting than after storage.

The core resistome determining the resistance genes which are not removed by any applied treatment, was also identified in the present study. These genes harbour resistance to tetracyclines (*tet36, tetA, tetQ, tetG, tetW, tetX, tetT, tetL, tetM*), aminoglycosides (*aadA5, aadA1, strB, aadA2, aadA9*), sulfonamides (*sul1, sul2*), trimethoprim (*dfrA1*), florfenicol (*floR*), antibiotics from MLSB group (*lnuB, mefA*) heavy metals (*merA*), as well as MGEs (*IntI1, intI2, tnpA, tnpA, IS1247*), and genes encoding the MDR phenotype (*qacEdelta1*). All of the listed genes harbour resistance to antibiotic groups commonly used in both veterinary and medical treatments (Zalewska and Popowska, 2020).

In general, following composting, livestock waste is mixed with different bulking materials to ensure optimal conditions for microbial growth and process development. Little is known of the type and chemical composition of the mixed biomass, and their effect on the quality and maturity of the compost produced. Careful mixture preparation is essential for balancing moisture, pH, and the C/N ratio for adequate aeration and microbial growth (Adhikari et al., 2009). When the physical structure is correct, the composting mixtures ensures oxygen for microorganisms, which is essential for mineralization; the oxygen availability mostly depends on the bulk density and moisture of the starting mixture (Agnew and Leonard, 2003; Trémier et al., 2009). The correct development of composting depends on its active and curing phase. Pezzola et al. (Pezzolla et al., 2021) demonstrated that the maximum attainable temperature, the time to reach it, and the pH and moisture depend on the used waste type and bulking agents. In addition, the dynamics of antibiotics removal appear to be closely influenced by the choice of bulking agent (sawdust rice husk, mushroom residues) (J. Zhang et al., 2019). Although these studies conclude that composting conditions affect the dynamics of antibiotic and ARG elimination from manure; no study has attempted to optimize the process. Studies have found treatment time and temperature to be the key factors affecting changes in ARG level during sludge treatment (Diehl and LaPara, 2010; Ma et al., 2011). In addition, water content and pH value appear to have a strong influence on antibiotic degradation (Wang and Yates, 2008); as changes in antibiotic levels also directly affect ARG levels [17], these two parameters also seem to have a strong influence on ARG profile. Furthermore, light seems to increase antibiotic and ARG removal thanks to the presence of natural photosensitizers in the environment, such as dissolved organic matter or porphyrins. When photosensitizers absorb the light, they produce singlet oxygen, which has strong oxidizing properties that can promote the conversion of antibiotics and disrupt ARGs (Engemann et al., 2008; Peak et al., 2007).

Studies on the factors determining ARG profiles during composting have found that certain physicochemical parameters, e.g., temperature, pH, ammonia content, and moisture, have a stronger effect on ARGs than the presence of host bacteria (Martinez, 2009; Qian et al., 2018; Zhang et al., 2018). While composting is a dynamic process influenced by the combined activity of a variety of bacteria, actinobacteria and fungi, the dynamics of the bacterial population are directly influenced by the environmental conditions including temperature, pH or moisture; for example, if a thermophilic species of bacteria carries the ARGs, their abundance may increase during the thermal phase (Duan et al., 2019).

## 3. Potential for horizontal gene transfer of ARGs

Horizontal gene transfer between different species has been recognized as a common and significant evolutionary process (Zhaxybayeva and Doolittle, 2011). It is most acutely demonstrated in the close interconnection between the resistome of manure-dwelling gut bacteria and potential pathogens, under constant selective pressure of the antimicrobial substances present in faeces. The main vessels of gene flow in microbial communities are plasmids which link distinct genetic pools (Halary et al., 2010). Our present findings indicate the presence of plasmids with a broad host range in both raw manure samples and in manure treated with composting or storage. Broad host range plasmids can replicate and stably maintain their gene payloads in taxonomically-distant species.

Our findings indicate the presence of plasmids from the Q, P, N, and W incompatibility groups (IncQ, IncP, IncN, and IncW). IncP plasmids may transfer between nearly all species of the Alpha-, Beta- and Gammaproteobacteria, and replicate in them. IncQ plasmids are broad host range plasmids that can proliferate in almost all Gram-negative bacteria. In addition, IncW and IncN plasmids can transfer and replicate in Gram-negative bacteria (Jain and Srivastava, 2013). The presence of mobile broad host range plasmids plays a key role in the dissemination of beneficial genetic traits, including resistance to antibiotics, metals, quaternary ammonium compounds and triphenylmethane dyes, and degradation of herbicides, both within the population of a certain species, and outside it (Jain and Srivastava, 2013; Zalewska et al., 2021).

Our study also identified *IncP_oriT* and *IncQ_oriT*; *oriT* is the origin of replication in Gram-negative bacteria transfer systems, and is crucial for transferring DNA from the donor to the recipient. Our samples were also found to include the *repA* gene, which encoded the initiation replication protein, ssDNA-dependent ATPase and DNA helicase, which most probably correlated with plasmid-copy number (Meyer, 2009), an internal gene necessary for transposition (*tnpA*), which is necessary for the recognition, cleavage, and integration of transposable elements (Ho, 2009), and a few insertion sequences (IS): IS614 (harboring ARGs, e.g., the *cfiA* gene) (Kato et al., 2003; Sóki et al., n.d.), IS1247 (encoding MDR phenotype – *aac(6’)IIc/ereA2/acc/arr/*ereA2) (Krauland et al., 2010) and IS1111 (harboring ARGs - *bla*_GES- 9_) (Poirel et al., 2005).

Our study compared the abundance of ARGs and MGEs in pig manure, both untreated and after storage or composting. The emergence and spread of ARGs are undoubtfully related to the presence of MGEs such as plasmids, transposases and integrases. A co-occurrence network constructed based on the ARGs and MGEs found in our samples confirmed the co-occurrence of MGEs, integrons and ARGs during both composting and storage. Although the co-occurrence network for the manure treated with storage has fewer nodes than that constructed during composting, it included more positive correlations between MGEs and ARGs, and these also tended to be more centralised: the largest cluster in the storage group includes 10 MGEs, compared to 7 MGEs in composting. Our findings clearly show that even during composting or storing, animal manure should be considered a hot spot for antibiotic resistance in which MGEs and ARGs are present together. Animal manure is a nutrient-rich environment inhabited by a highly-dense bacterial population, which with the addition of antibiotics and heavy metals, makes favourable conditions for HGT. It has been proven that the presence of antibiotics in manure increases the activity of transposases, thus resulting in more frequent excision or integration of gene cassettes in integrons (Heuer et al., 2011).

The *tet* genes, which were found to be the most prevalent genes in the untreated pig manure, are often identified as plasmid components. Their spread is further facilitated by insertion into transposons such as Tn6298, Tn1721, Tn6303, or IS1216. For example, the *tetW, tetQ, tetQ* genes are known to be common in the host gastrointestinal microbiota, indicating that the genes demonstrate high stability within the population (Heuer et al., 2011; Leclercq et al., 2016). To date, three sulfonamide-resistance genes have been identified: *sul1*, frequently found within class 1 integrons, *sul2*, found on a broad-host-range plasmid in *E. coli*, and *sul3*, also found together with class 1 integrons in a variety of environments. Due to their location on MGEs, together with integrons and transposons, sulfonamide-resistance genes have high transfer potential and can be found in many bacterial species (Lima et al., 2020).

Pig manure is also relatively abundant in beta-lactam resistance genes, since they commonly receive large levels of antibiotics belonging to these groups. The genes are often located within IncN plasmids (Binh et al., 2008). IncN plasmids may carry *bla_CTX-M_*, suggesting that they play a crucial role in the dissemination of ESBL strains in manure. Moreover, the genes coding for the ESBL phenotype are often located on IS*Ecp1* or IS*CR1* (Lee et al., 2020). Macrolide-resistance genes (e.g., *ermB*) and tetracycline-resistance genes may also be located on MGE, such as the Tn*916*-Tn*1545* family of conjugative transposons. Other interesting genes also have been found in manure associated with MGEs, e.g., the colistin-resistance gene has been found on diverse plasmid replicon types, such as IncX4, IncI, IncFII, IncX1, and IncQ1, in *E. coli* (Lima et al., 2020). Other reports have noted the presence of *ampC*, *qnr*, carbapenemase-resistance genes originating from animal manure on plasmids with a broad host range, such as replicon type plasmids (IncP-1, IncQ, IncN, incW, or IncF), underlining their high transfer potential (Chee-Sanford et al., 2009).

## 5. Conclusions

Antibiotics are a critical tool for fighting bacterial infections in humans and animals. However, there is an increasing need for greater care in their use to mitigate the development of widespread resistance. For example, it is advisable to treat only animals with clinical manifestations instead of employing prophylactic or metaprophylactic treatment for an entire herd. Numerous reports indicate that the use of animal fertilizers also entails considerable risk of spreading antibiotic resistance. Pig manure, with its abundant and diverse ARGs, being a major source of resistance genes, poses a particular threat to public health. Microbes from manure, compost, or soil containing the ARGs are subject to dispersal via runoff into rivers, leaching to subsurface waters, air dispersal via dust, human travel, and distribution of agricultural products, including compost for gardening; such processes may expand local contamination to regional or even global scale.

The present study examined the effect of two treatments, storage and composting, on the potential of pig manure from a medium-sized livestock farm to harbour antibiotic resistance. The study was performed on the same manure samples, at the same time and under controlled conditions. Furthermore, both sets of samples were subjected to the same analyses of microbial and resistome composition as well as their physicochemical properties.

Our findings reveal an alarming diversity and abundance of ARGs, indicating that raw pig manure represents a ubiquitous reservoir of ARGs and MGEs, which poses a considerable risk of ARGs spreading into the environment when released. The co-occurrence of AGRs and MGEs increases the risk of transfer of ARGs from bacteria inhabiting livestock animals and their environments to human-associated bacterial pathogens, resulting in further spread among human populations. The presence of such resistant strains limits the choice of possible treatment strategies in infection.

Generally, positive correlations were observed between microbial population composition and the presence of specific ARGs and MGEs and genes coding the MDR mechanism, and between ARGs and MGEs in all tested systems. It was found that composting and storage resulted in differential changes in the quantities of ARG groups, while others demonstrated lower, or even higher, gene copy numbers; however, MGEs were found to be one of the groups whose abundance increased during both processes.

Our co-occurrence network encompassing both MGEs and ARGs imply that horizontal transfer of genes may occur not only in raw manure, but also during composting or storage before field application. Although the relative abundance of MGEs generally increases during both treatments and their prevalence is associated with a high diversity of resistance genes, a lower overall abundance of MGEs is observed during composting than during storage; this suggests that although composting may not be sufficient to completely limit the spread of ARGs, it is more efficient than simple storage.

Hence, it can be concluded that the use of composted manure is a safe strategy to ensure soil fertility and high humus content. It represents a promising alternative to mineral fertilizers, thus contributing to a more ecological way of farming crops.

## CRediT authorship contribution statement

**MZ -** Methodology, Formal analysis, Investigation, Data Curation, Writing - Original Draft, Visualization; **AB** - Formal analysis, Investigation; **AC** - Investigation; **MP** - Conceptualization, Writing - Review & Editing, Supervision, Project administration, Funding acquisition

## Acknowledgments

The research was funded by National Science Centre, Poland (2017/25/Z/NZ7/03026), grant under the European Horizon 2020, in the frame of the JPI-EC-AMR Joint Transnational Call (JPIAMR), JPI-EC-AMR JTC 2017, project INART – “Intervention of antibiotic resistance transfer into the food chain” to MP and partially in the frame of the “Excellence Initiative – Research University (2020-2026)” Program at the University of Warsaw.

---

**Table S1.** Physicochemical parameters of pig manure: raw and after composting and storage

| Parameters | Sample type |  |  |  |
| --- | --- | --- | --- | --- |
|  | Control | Storage (4M) | Compost (10W) |  |
| pH | 7.0 | 7.2 | 7.4 |  |
| DM<br>(% of w.t.) | 27.63±1.28 | 32±1.02 | 54.25±0.11 |  |
| OM<br>(% of d.m.) | 89.60±5.17 | 91.45±1.47 | 118.01±0.89 |  |
| Manurial<br>[% d.m.] |  |  |  |  |
| (N-NH<sub>4</sub><sup>+</sup>) | 1.25±0.09 | 1.18±0.10 | 2.13±0.12 |  |
| Total N | 3.11±0.23 | 6.12±0.31 | 6.85±0.09 |  |
| Total P | 1.55±0.15 | 1.14±0.08 | 0.81±0.02 |  |
| Total Mg | 1.28±0.11 | 1.12±0.07 | 0.88±0.02 |  |
| Total Ca | 3.62±0.31 | 2.86±0.26 | 2.32±0.08 |  |
| Heavy metals<br>(mg/kg) |  |  |  | Permissible standard for agricultural land (mg/kg)* |
| Chrome (Cr) | 4.37±0.31 | 4.42±0.24 | 4.65±0.29 | 150-500 |
| Copper (Cu) | 128±9.00 | 146±8.00 | 215±11.00 | 100-300 |
| Cadmium (Cd) | 0.561±0.02 | 0.612±0.02 | 0.624±0.01 | 2-5 |
| Nickel (Ni) | 6.41±0.56 | 6.88±0.85 | 7.39±0.48 | 100-300 |
| Lead (Pb) | 2.49±0.17 | 2.52±0.11 | 2.68±0.12 | 100-500 |
| Zinc (Zn) | 796±15.00 | 809±14.00 | 816±13.00 | 300-1000 |
| Mercury (Hg) | 0.0468±0.01 | 0.0532±0.01 | 0.0571±0.01 | 2-5 |

\*Values depending on the type/properties of arable soil in accordance with applicable standards in Poland
DM – dry matter; OM – organic matter

**Table S2.**
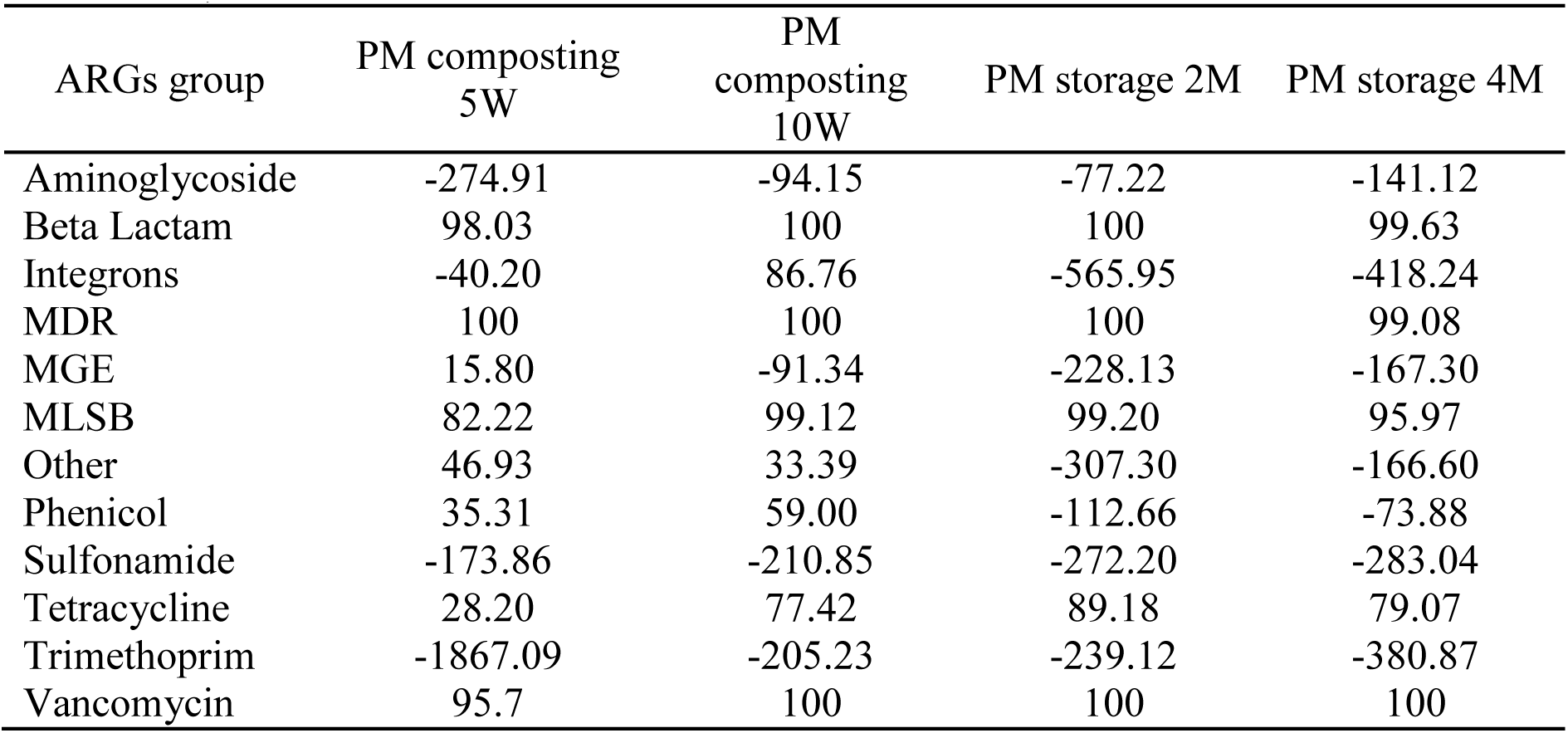
Reduction [%] of the relative abundances of ARG groups during composting (PM composting 5W, PM composting 10W; composted samples after five weeks and ten weeks, respectively), and stored (PM stored 2M, PM stored 4M; stored samples after two months and four months)

**Table S3.**
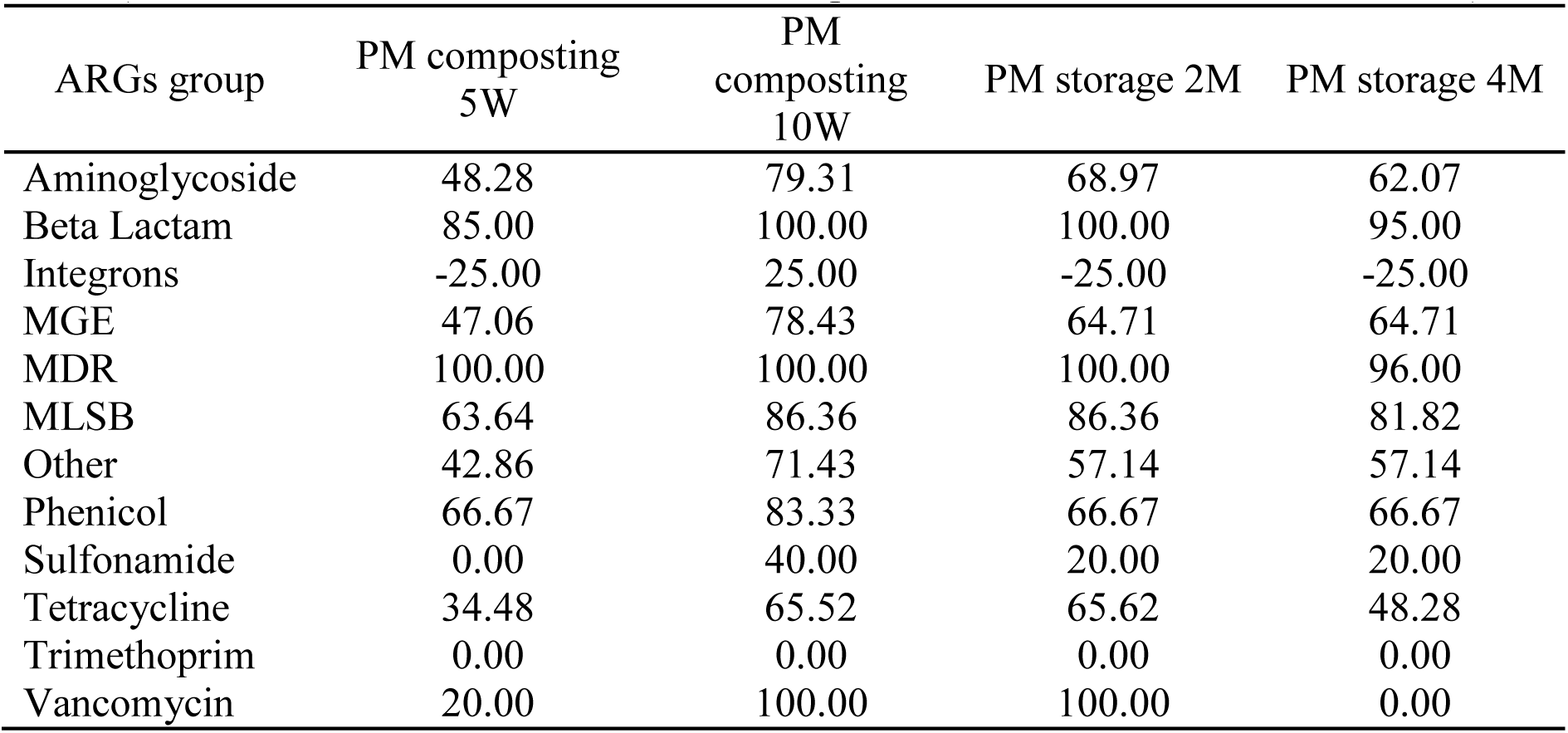
Reduction [%] in ARG level during composting and storage (PM composting 5W, PM composting 10W; composted samples after five weeks and ten weeks, respectively), and stored (PM stored 2M, PM stored 4M; stored samples after two months and four months)

